# Brain-wide representations of behavior spanning multiple timescales and states in *C. elegans*

**DOI:** 10.1101/2022.11.11.516186

**Authors:** Adam A. Atanas, Jungsoo Kim, Ziyu Wang, Eric Bueno, McCoy Becker, Di Kang, Jungyeon Park, Cassi Estrem, Talya S. Kramer, Saba Baskoylu, Vikash K. Mansingkha, Steven W. Flavell

**Affiliations:** Picower Institute for Learning & Memory, Massachusetts Institute of Technology, Cambridge, MA, USA; Computational and Systems Biology Program, Massachusetts Institute of Technology, Cambridge, MA, USA; Department of Brain & Cognitive Sciences, Massachusetts Institute of Technology, Cambridge, MA, USA; Department of Electrical Engineering and Computer Science, Massachusetts Institute of Technology, Cambridge, MA, USA; MIT Biology Graduate Program, Massachusetts Institute of Technology, Cambridge, MA, USA

## Abstract

Changes in an animal’s behavior and internal state are accompanied by widespread changes in activity across its brain. However, how neurons across the brain encode behavior and how this is impacted by state is poorly understood. We recorded brain-wide activity and the diverse motor programs of freely-moving *C. elegans* and built probabilistic models that explain how each neuron encodes quantitative features of the animal’s behavior. By determining the identities of the recorded neurons, we created, for the first time, an atlas of how the defined neuron classes in the *C. elegans* connectome encode behavior. Many neuron classes have conjunctive representations of multiple behaviors. Moreover, while many neurons encode current motor actions, others encode recent actions. Changes in behavioral state are accompanied by widespread changes in how neurons encode behavior, and we identify these flexible nodes in the connectome. Our results provide a global map of how the cell types across an animal’s brain encode its behavior.

## INTRODUCTION

Animals can generate a vast array of behavioral outputs that vary depending on their environment, context, and internal state. The neural circuits that control these behaviors are distributed across the brain, and their dynamic interactions underlie the neural control of behavior. To decipher how these circuits work, it will be critical to relate the activity of this full population of neurons to specific features of animal behavior. However, it is immensely challenging to measure brain-wide activity and comprehensive behavioral information of a freely-moving animal. For this reason, it has remained unclear how neurons and circuits across entire nervous systems represent an animal’s diverse behavioral repertoire, and how this flexibly changes depending on an animal’s context or internal state.

Recent studies suggest that internal states and moment-by-moment behaviors are associated with widespread changes in neural activity spanning many brain regions (Allen et al., 2019; Brezovec et al., 2022; Hallinen et al., 2021; Marques et al., 2020; Musall et al., 2019; Schaffer et al., 2021; Stringer et al., 2019). For example, behavioral states, such as active versus quiet wakefulness, and homeostatic states, like thirst, are associated with changes in neural activity in many brain regions (Allen et al., 2019; Niell and Stryker, 2010; Stringer et al., 2019). In addition, instantaneous motor actions are associated with altered neural activity across a surprisingly large number of brain regions (Musall et al., 2019; Stringer et al., 2019). This gives rise to a view that there are ongoing representations of an animal’s behavior and its state in many brain regions. However, our understanding of how global neural dynamics spanning many brain regions encodes behavior remains limited. For instance, in mammalian systems, representations of motor actions occur throughout the brain: in the cortex, cerebellum, midbrain, spinal cord, and more. But the actual forms of the neural representations – how the neurons and circuits encode a diverse set of motor outputs – in most of these regions are still unknown. In addition, given the vast number of cell types involved and their broad spatial distributions, characterizing this entire system is not yet tractable.

Adult hermaphrodites of the nematode *C. elegans* have a compact nervous system consisting of 302 defined neurons with known connectivity (Cook et al., 2019; White et al., 1986; Witvliet et al., 2021). *C. elegans* generates a well-defined repertoire of motor programs: locomotion, feeding, head oscillations, defecation, egg-laying, and postural changes. Previous studies of *C. elegans* behavior have shown that this animal’s nervous system is subject to modulation, such that animals can express different behaviors as they switch between different behavioral states (Flavell et al., 2020). For example, animals enter sleep-like states during development and after intense stress (Raizen et al., 2008; Van Buskirk and Sternberg, 2007). Awake animals exhibit different locomotion patterns during different foraging states, like roaming versus dwelling (Flavell et al., 2013; Fujiwara et al., 2002; Ji et al., 2021). In addition, sudden aversive stimuli induce long-lasting behavioral states in which animals’ arousal increases for minutes after the initiating stimulus (Ardiel et al., 2017; Chew et al., 2018). Thus, *C. elegans* provides a system where it may be feasible to comprehensively describe how behavioral variables are encoded by activity across an entire nervous system, and how this can flexibly change over time.

Recordings of *C. elegans* neurons in freely-moving animals have identified some individual neurons that reliably encode specific behavioral features. The neurons AVA, AIB, and RIM encode backwards motion; AVB, RIB, AIY and RID encode forwards motion; SMD encodes head curvature; and HSN encodes egg-laying (Gordus et al., 2015; Kaplan et al., 2020; Kato et al., 2015; Li et al., 2014; Lim et al., 2016; Luo et al., 2014; Roberts et al., 2016; Zhang et al., 2008). Brain-wide calcium imaging in immobilized animals has identified population activity patterns associated with fictive locomotion dynamics (forward/reverse/turn) (Kaplan et al., 2020; Kato et al., 2015). Indeed, velocity and curvature can be decoded from population activity in moving animals (Hallinen et al., 2021), suggesting that this information is broadly reflected in neural activity. However, there is still a major gap in our understanding of how the vast majority of the neurons in the *C. elegans* brain encode features of the animal’s behavior as it moves freely. This is due in large part to the technical difficulty of recording comprehensive, high-signal-to-noise (SNR) neural/behavioral datasets that would permit such an understanding. Thus, the main modes of behavior representation across the nervous system – timescales of representation, how the diverse motor programs are conjunctively encoded, and state-dependent changes in representation – remain unknown.

Here, we elucidate how neurons across the *C. elegans* brain encode the animal’s behavior. We developed technologies that allowed us to simultaneously record high-fidelity brain-wide activity and the diverse motor programs of 37 freely-moving animals. We then devised a generalizable probabilistic encoding model that can fit most recorded neurons, providing an interpretable description of how each neuron encodes behavior. By also determining neural identity in 12 of these brain-wide recordings, we created an atlas of how most of the *C. elegans* neuron classes encode behavior. We find that many neuron classes have combined representations of multiple motor programs. Moreover, while many neuron classes represent current motor actions, others also represent recent motor actions, such that past behavior can be decoded from current population activity. Finally, we show that ~20% of the neurons can flexibly change how they encode behavior over time and that behavioral state changes are associated with this type of remapping, revealing a striking degree of flexibility in this system. Our results provide one of the first views of how activity across the defined cell types of an animal’s brain encodes quantitative features of its diverse motor outputs.

## RESULTS

### Technologies to record brain-wide activity and a diverse set of motor programs

To determine how neurons across the *C. elegans* brain encode features of the animal’s behavior, we developed a new microscopy platform for brain-wide calcium imaging in freely-moving animals and wrote new software to fully automate processing of these recordings. We constructed a transgenic *C. elegans* strain that expresses NLS-GCaMP7f (a calcium sensor) and NLS-mNeptune2.5 (a red fluorescent protein) in all neurons. Recording nuclear-localized GCaMP makes it feasible to record brain-wide activity, though this approach will miss local compartmentalized calcium signals in neurites (Hendricks et al., 2012). After integration of the transgenes into the animal’s genome, we confirmed that the transgenic animals’ behavior was phenotypically normal, using assays for chemotaxis and associative learning (Fig. S1A). Animals were recorded on a custom microscope with two light paths, inspired by recent work (Fig. 1A-C; Nguyen et al., 2016; Venkatachalam et al., 2016). The lower light path is coupled to a spinning disk confocal for volumetric imaging of fluorescent signals in the head. The upper light path has a low-magnification objective and near-infrared (NIR) brightfield configuration to capture images of the worm for behavior quantification (Movie S1). To allow for closed-loop animal tracking, the location of the worm’s head is identified in real time (at 40 Hz) with a deep neural network (Mathis et al., 2018) and input into a PID controller that automatically moves the microscope stage to keep the animal centered. This permits us to record brain-wide calcium signals and behavior in a freely-moving animal.

**Figure 1.**
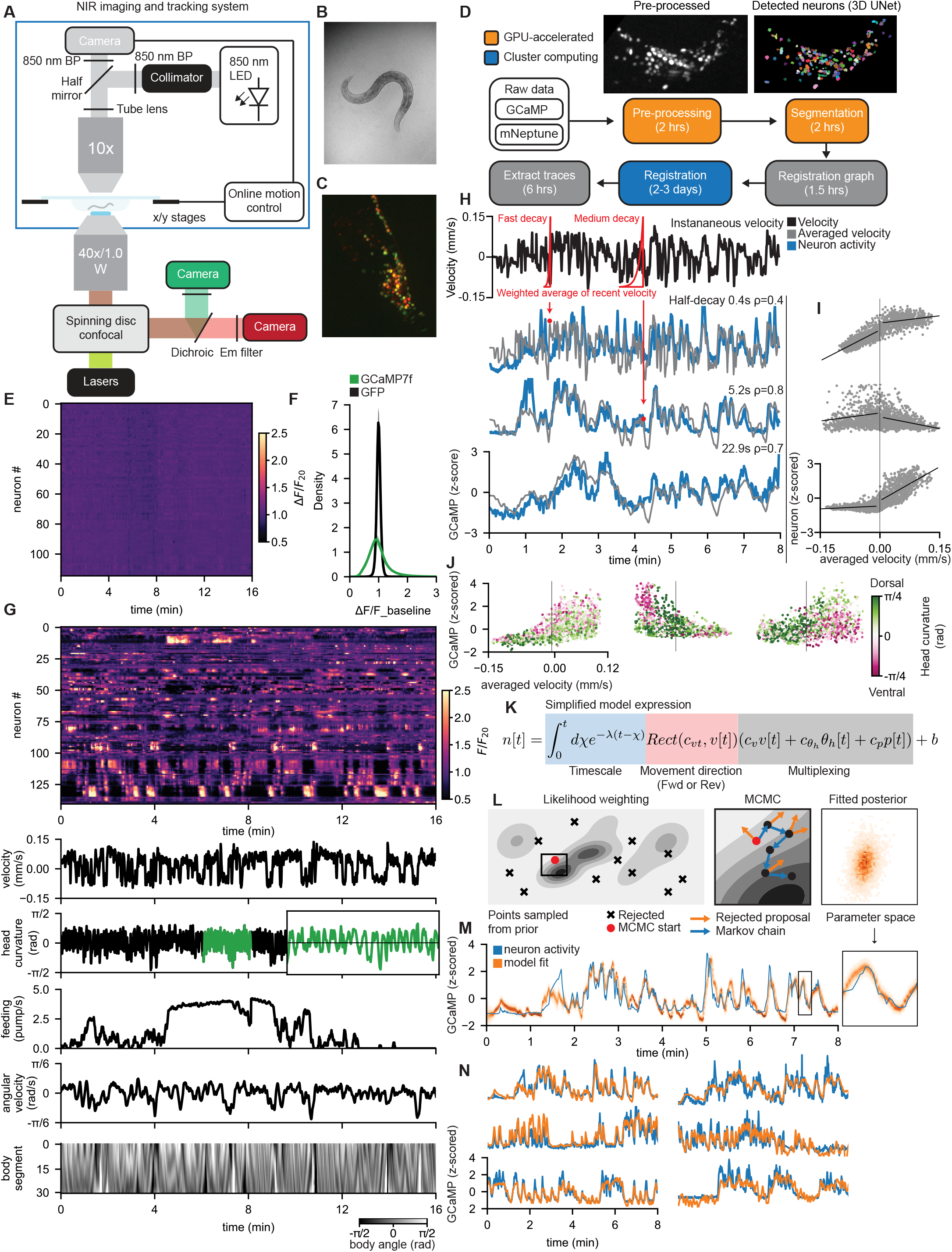
The *C. elegans* Probabilistic Neural Encoding Model (CePNEM) can reveal how neurons across the *C. elegans* brain represent behavior. (A) Light path of the microscopy setup. On the upper light path (“NIR imaging and tracking system”), the 850-nm (NIR) LED is collimated and filtered (850-nm bandpass). The illumination light is reflected downward by the half mirror into the 10x objective, illuminating the sample. The reflected light is collected by the same objective and passed through the half mirror. The image is filtered (850-nm bandpass) and captured by the camera. The captured image (panel B) is processed by the online tracking system, which sends out commands to the stage to cancel out the motion. On the bottom light path, the spinning disc confocal setup illuminates and collects the fluorescence signal from the sample. The collected signal is split by a dichromatic mirror and captured by two cameras. (B) Example image of a worm collected through the NIR brightfield light path. (C) Example image of a confocal volume (maximum intensity projection) captured at the same time as in (B). (D) Automatic Neuron Tracking System for Unconstrained Nematodes (ANTSUN) software pipeline to process and extract GCaMP signals from the confocal volumes over time. Detailed descriptions of each of the steps depicted in this cartoon are provided in Methods. (E) F/F_20_ (F_20_ is the 20^th^ percentile of F across the time series for the neuron) heatmap of neural traces collected from a pan-neuronal GFP control animal. Data are shown using same color scale as GCaMP data in (G). (F) Comparison of variation in F/F_20_ from the extracted traces in all neurons in 3 GFP control animals (σ = 0.074) and 14 GCaMP animals (σ=0.392). (G) A full example dataset, showing a F/F_20_ (F_20_ is the 20^th^ percentile of F across the time series for the neuron) heatmap of neural traces together with the animal’s velocity, head curvature, feeding behavior, angular velocity (change in animal’s heading over time), and body angles (a vector of angles from head to tail that define the shape of the animal). For head curvature, an inset (green) shows a zoomed in region of the behavioral trace to illustrate fast head oscillations. (H) Three example neurons from one animal that encode velocity over different timescales. Each neuron (blue) is correlated with an exponentially-weighted moving average (red) of that animal’s recent velocity, but over different timescales (gray traces). Note that the three gray traces show velocity from the same animal over the same time segment, but convolved with exponential decay kernels that have different half-decay times, as is illustrated by the red kernels overlaying the instantaneously velocity trace from that animal (top). The different example neurons’ correlation coefficients *ρ* to these filtered velocity data are displayed as insets; we also display the exact half-decay times of the exponential filters used. (I) Example tuning (velocity vs neural activity) scatterplots for three example neurons (different from those in H) showing how their activity relates to velocity (see Methods). The dots are individual timepoints (each with a neural activity and corresponding behavioral measurement) for these three example neurons. Altogether, the full set of dots reveal how each neuron’s activity changed as a function of velocity. Separate trendlines were fit to all datapoints for reverse and forward velocity. (J) Example tuning scatterplots for three example neurons displaying how neurons’ activities can combine information about the animal’s head curvature (color) and velocity (x axis). The dots are individual timepoints (each with a neural activity and corresponding behavioral measurement) for these three example neurons. Head curvature at each timepoint is indicated by the color of the dot. Altogether, the full set of dots reveal how each neuron’s activity changed as a function of velocity and head curvature. Note that for each neuron the red and green dots separate from one another only for negative or positive velocity values (corresponding to reverse or forward movement, respectively). This indicates that the neurons vary their activity based on head curvature only during forward (neurons on left and right) or reverse (neuron in middle) movement. (K) Simplified expression of the deterministic component of CePNEM. This model is applied to each neuron in the brain-wide recordings. Each neuron is modeled as the recent weighted average of multiple behavioral predictor terms. Note that in this simplified depiction of the model Equation 1 in the text. We represent the effect of timescale via an integral with parameter *λ*, whereas Equation 1 represents timescale via recursion with parameter s, which we then transform into and report as a half-decay time *τ*_1/2_. (L) Left and Middle: Schematic demonstrating how the MCMC fitting process is initialized and fit. Likelihood weighting selects a particle with the best fit to the neural and behavioral data. An MCMC process is then used to determine the posterior distribution. Gray shading here indicates model likelihood given the parameters in that region of parameter space. Right: an example posterior distribution that results from the fitting process being run on a neural trace. This is shown for just two of the model’s parameters (x- and y-axes here) for this illustrative purpose. (M) An example neuron trace (blue) that was fit with CePNEM. This fit resulted in a posterior distribution of model parameters. A model trace *n*[*t*] was generated from each set of parameters drawn from the posterior, and a heatmap of all such models is plotted in orange. (N) Example neural traces overlaid with the median of all posterior CePNEM fits for that neuron (referred to henceforth as median CePNEM fits).

We wrote a software suite (Automatic Neuron Tracking System for Unconstrained Nematodes, or ANTSUN) to automatically extract calcium traces from these videos (Fig. 1D). In this software package, we used the time-invariant mNeptune2.5 signal to determine the locations of neurons and register images from different timepoints to one another. First, a custom 3D U-Net (Wolny et al., 2020) was used to locate and segment all neurons in all timepoints. Second, we constructed a registration graph based on posture similarity, in which timepoints were represented as nodes and they were connected by an edge if the postures at the two timepoints were sufficiently similar to make volume registration tractable. Third, we solved all volume registration problems in the graph. Fourth, we devised a distance metric that indicates the likelihood that any two neurons recorded at different timepoints were the same. We then used a custom clustering approach to link neurons’ identities over time (see Methods). Finally, fluorescence (F) was computed as the ratio of GCaMP to mNeptune intensity at each time point. To ensure that our approach accurately tracked the same neuron over time as animals moved, we recorded a control strain expressing NLS-GFP at different levels in different neurons (*eat-4::NLS-GFP*), along with pan-neuronal NLS-mNeptune2.5 (Fig. S1B). Mistakes in linking neurons’ identities over time would be obvious in this strain, since GFP levels would fluctuate in a neural trace if timepoints were sampled from different ground-truth neurons. We quantified the prevalence of errors of this type, taking into account the variance in GFP signal between neurons, and found that neural traces were correctly sampled from individual neurons in 99.7% of their recorded frames. Thus, neural identification errors are negligible in these datasets. We also estimated the degree of motion artifacts in our data by recording a transgenic strain expressing pan-neuronal NLS-GFP and NLS-mNeptune2.5 (Fig. 1E, compare to Fig. 1G; see also Fig. S1C). GFP should be constant over time, so any signal fluctuations would be due to motion or image processing artifacts. We found that the distribution of fluorescent signals over time was far more narrowly distributed for GFP, compared to GCaMP7f, suggesting that motion artifacts are also negligible (Fig. 1F; standard deviations of GCaMP and GFP distributions were 0.392 and 0.074, respectively). Nevertheless, we used the GFP datasets to correct and control for any such artifacts in all analyses below (see Methods).

We also wrote software that extracts a diverse list of behavioral variables from the NIR brightfield images. In each frame, the animal is detected via a convolutional neural network and a spline is fit to its centerline. Velocity is computed as the rate of movement of the animal’s head projected onto the direction the animal is facing. Angles along the centerline parameterize the worm’s head and body posture. Feeding (or, pharyngeal pumping) is manually quantified from videos played at 25% of real-time speed. From these variables, we derive additional behavioral features: movement direction (forward/reverse), angular velocity, head curvature (oscillatory bending of the head), and more. We examined the data closely for egg-laying and defecation events, but found that animals did not exhibit these behaviors under the recording conditions, so they are not included in any of the analyses below. Altogether, these advances permit us to quantify brain-wide calcium signals and a diverse list of behavioral variables from freely-moving *C. elegans*.

### A probabilistic neural encoding model reveals how each *C. elegans* neuron encodes quantitative features of the animal’s behavior

To determine how neurons across the *C. elegans* brain encode the animal’s behavior, we recorded brain-wide activity and corresponding behavioral data from 14 animals as they freely explored a sparse food environment. Each recording was ~16 minutes long and we obtained data from 143 ± 12 head neurons per animal (example in Fig. 1G; Movie S1). Our objective was to precisely describe how each neuron “encodes” or “represents” the animal’s behavior, in other words how its activity is quantitatively associated with features of the animal’s behavior. Such an association could be due to a given neuron causally influencing behavior or, alternatively, receiving proprioceptive or corollary discharge signals relevant to behavior; both of these types of representations are essential for a nervous system to properly control behavior. Our initial efforts to build models of how neurons encode behavior revealed three important features of how neural activity relates to behavior that were not fully characterized in prior work. We describe these features here and provide a systematic identification of all neurons with these features below (including summary statistics across all neurons and animals; see Fig. 2).

**Figure 2.**
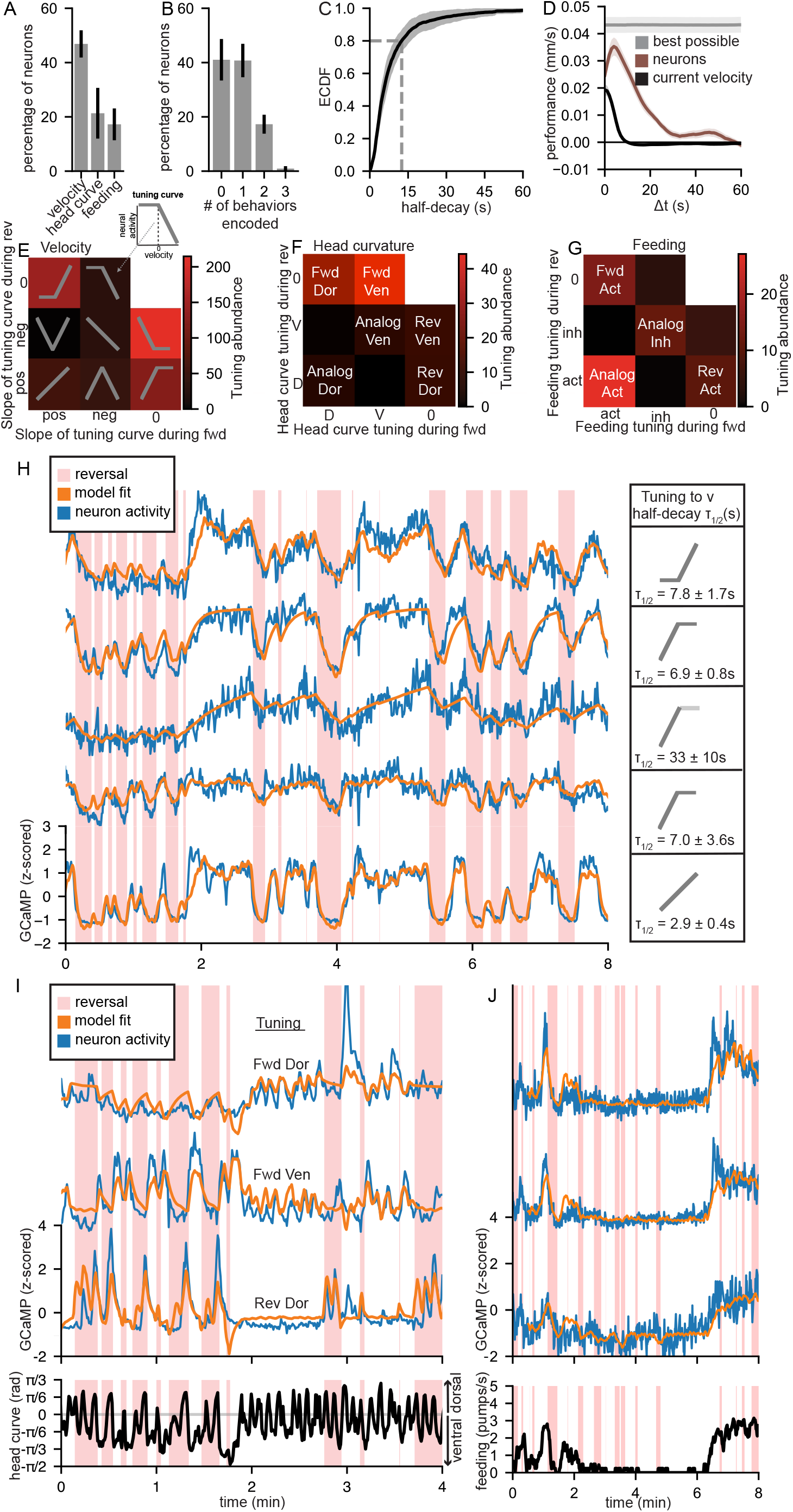
Rich and varied representations of behavior across the *C. elegans* brain, spanning multiple timescales. (A) Mean fraction of all detected neurons in the brain that encode velocity, head curvature, and feeding in 10 animals (datasets with insufficient variance in pumping behavior are excluded from this particular analysis - see Methods). Error bars are the standard deviation between animals. (B) Mean fraction of all detected neurons in the brain that encode 0, 1, 2, or 3 of the behaviors in the model (velocity, head curvature, or feeding) in the same 10 animals as in (A). Error bars are the standard deviation between animals. (C) Mean ECDF of the median model half-decay time of all neurons demonstrated to encode at least one behavior in 14 animals. The shaded region represents the standard deviation between animals. (D) Performance of linear decoders trained to predict velocity at times in the past (x-axis) from current neural activity (red). Performance was defined as the difference in error (computed as the standard deviation of the difference between actual velocity and predicted velocity, in mm/sec) between the actual decoders and control decoders that used time-shifted values of the predictive variables (see Methods for more details). The velocity values to be predicted were each averaged over a 10 second sliding window centered Δt seconds into the past. All neurons with significant encoding of velocity were used as predictor terms. For comparison, a decoder was also trained to make this prediction based on current velocity (black), so that we could estimate the degree to which current velocity predicts past velocity via autocorrelation. Another “best possible” decoder was also trained to make this prediction based on current and past velocity (gray), which should be able to perform nearly-perfectly (since it is given the information it is trying to predict) and thus estimate the best possible performance of such a decoder. Note that the use of the 10-second sliding window causes the current velocity decoder to underperform the best possible decoder even at Δt = 0. The observation that neurons predict velocity better in the past (Δt ≈ 5) than present likely relates to the fact most velocity-encoding neurons in the head represent recent, rather than instantaneous, velocity. Error shading indicates standard deviation across animals. (E) A categorization of how the full set of velocity-encoding neurons represent velocity. Based on each neuron’s tuning to velocity during forward and reverse movement (i.e. the slope of its tuning curve for velocities >0 and <0, respectively), it could potentially be categorized into one of eight groups. All velocity encoding neurons were sorted into these categories based on a statistical analysis of their CePNEM fits. The overlaying gray traces indicate the prototypical tuning curve for each category (inset in the upper right illustrates how to interpret the gray tuning curves, using an example). See Methods for more details. Color reflects “tuning abundance,” which is the number of detected neurons in each bin, scaled by their velocity encoding strength. (F) Same as (E), but for head curvature encoding neurons. (G) Same as (E), but for feeding encoding neurons. (H) Five example neurons from the same animal that all encode forward locomotion, together with CePNEM-derived tuning curve diagrams for each neuron, and the mean and standard deviation each neuron’s half-decay time *τ*_1/2_. Note that although all five neurons encode forwards velocity, their neural activity traces have notable differences in dynamics. Importantly, the model (orange) captures these differences. The third neuron lacked statistical significance on the velocity slope variable during forward locomotion, so the tuning curve is in a lighter shade to reflect this uncertainty. (I) Three example neurons from the same animal that all encode head curvature in conjunction with movement direction, together with CePNEM-derived tuning parameters for each neuron (shown above each neural trace). Neural traces are shown in blue and median model fits are shown in orange. Head curvature of the animal is shown below. (J) Three example neurons from the same animal that all encode feeding information. All three neurons have the “Analog Act” tuning. Neural traces are shown in blue and median model fits are shown in orange. Feeding rate of the animal is shown below.

First, while examining neurons that were more active during forward or reverse velocity, we found that these neurons encode behavior over a surprisingly wide range of timescales, which had not been described before. We observed that the activity of individual neurons that encode velocity was precisely correlated with a weighted average of the animal’s recent velocity, in some cases stretching back in time as far as 30sec. Specifically, the neurons were strongly correlated with an exponentially weighted average of recent velocity. The decays of the exponentials, which determine how much a given neuron’s activity weighs past versus present velocity, varied widely across neurons (range of half-decay (τ_1/2_): 0.9 – 31.7 sec; the half-decay of the GCaMP7f sensor in live neurons is <1 sec, and estimates of cell type variability in its decay are <2-fold; (Dana et al., 2019; Wei et al., 2020)). Fig. 1H illustrates this by showing individual neuron traces from an example animal, along with its velocity that has been convolved with exponential filters with varying decay times. Individual neurons that are strongly correlated with velocity integrated over each of these specific timescales are also shown. We also observed a broad range of timescales for neurons that encode other behavioral features (see below). These data suggest that the neurons that encode *C. elegans* behavior differ in how much they reflect the animal’s past versus present behavior.

Second, we observed that neurons could reflect individual behaviors in a more heterogeneous fashion than expected. Focusing on velocity specifically, each neuron’s representation of velocity can be captured by a tuning curve that relates the neuron’s activity to velocity. The shapes of these tuning curves were quite different for different neurons. Some neurons displayed analog tuning, where their activity changed monotonically from fast reverse to fast forward movement (i.e. the slope of the tuning curve was the same across all velocity values). However, other neurons displayed evident “rectification”, in which the slopes of their tuning curves during reverse and forward velocity differed (Fig. 1I). Finally, many velocity-encoding neurons could not be classified as “forward” or “reverse,” but instead displayed other tunings, for example encoding slow locomotion regardless of movement direction (Fig. 1I, middle). These data suggest that neurons that encode velocity can exhibit different activity profiles, reflecting overall speed, movement direction, or finely tuned aspects of an animal’s forward or reverse movement.

Third, we found that many neurons conjunctively represent multiple distinct motor programs. For example, most neurons whose activities were correlated with the oscillatory bending of the worm’s head showed different tunings to head curvature during forwards versus reverse movement. In fact, many neurons represented information about head curvature only during either forward or reverse movement. Fig. 1J shows example tuning curves of these neurons to velocity, but the datapoints are also colored based on the animal’s head curvature, which reveals the joint tuning to both motor programs (note that green and red dots are separated from each other only during positive or negative velocity values, depending on the neuron). Similarly, many neurons conjunctively represented the animal’s velocity and feeding rate. These data suggest that a considerable number of *C. elegans* neurons encode multiple motor programs in combination, commonly referred to as “multiplexing” or “mixed selectivity.”

Based on these observations, we next sought to construct a computational model that could be used to reveal how each neuron encodes features of the animal’s behavior. Our approach was to construct an encoding model that uses behavioral features to predict each neuron’s activity (Equation 1; see Methods for details). The form of our model was generalizable, meaning that it could be applied to any neuron in our recordings to reveal how it encodes behavior. Each neuron’s activity was modeled as an exponentially weighted average of the animal’s recent behavior with a single fit decay parameter *s* for each neuron, allowing for different timescale encoding. Neurons can additively weigh multiple behavioral predictor terms (based on the coefficients *c_v_, c_θh_*, and *c_p_* for velocity, head curvature, and pumping/feeding, respectively), which can each multiplicatively interact with the animal’s movement direction parameterized by *c_vT_*. This allows for heterogeneous rectified and non-rectified tunings to velocity and other behavioral features, as well as multiplexing. To determine whether each of the model parameters were necessary to explain neural activity, we compared the goodness of fit of the full model to models with one parameter deleted (and to a fully linear model), and found that deletion of any parameter significantly increased the error of the model fits (Fig. S2A). We also fit more complex models (for example, a model where all behavioral predictors can multiplicatively interact with one another or models with rectification terms for other behavioral parameters), but found that this did not improve model performance.

The parameters of the model are interpretable, so the model fits allow us to describe how each neuron encodes each behavioral feature. However, because the model is fit on a finite amount of neural/behavior data, these parameters have a level of uncertainty that is important to estimate. For this reason, when fitting the model for each neuron, we determined the posterior distribution of all model parameters that were consistent with our recorded data, where consistency was defined as likelihood in the context of a Gaussian process noise model parameterized by *σ_noise_, σ_SE_*, and *ℓ* (see Methods). This allowed us to quantify our uncertainty in each model parameter and perform meaningful statistical analyses. The posterior distribution was determined using the probabilistic programming language Gen (Fig. 1L-M; Cusumano-Towner et al., 2019). We used a procedure where 100,000 particles randomly positioned in parameter space were filtered by likelihood weighting, after which a Markov Chain Monte Carlo (MCMC) process on the best particle was used to determine the posterior distribution of model parameters (Fig. 1L-M; see Methods for additional details, illustrating distinctive features of Gen that are not available in widely used probabilistic programming languages such as Stan). We confirmed the validity of this approach using simulation-based calibration, a well-known technique in computational Bayesian statistics for ensuring that MCMC approximations are sufficiently accurate (Fig. S2B; Talts et al., 2020).

### Equation 1: The C. elegans Probabilistic Neural Encoding Model (CePNEM) expression

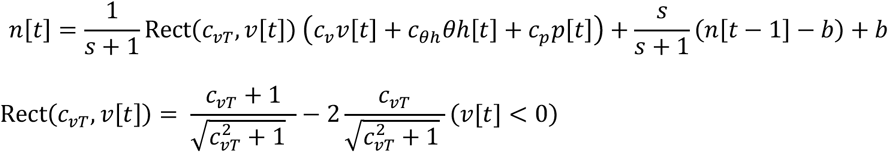

Observed neural activity 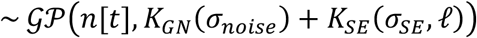

| Parameter | Meaning |
| --- | --- |
| Observed neural activity | Observed neural activity trace (z-scored) |
| $v[t]$ | Observed worm velocity |
| $\theta h[t]$ | Observed worm head curvature |
| $p[t]$ | Observed worm pumping rate |
| $n[t]$ | Modeled neural activity |
| $\text{Rect}(c_{vT}, v[t])$ | Locomotion direction rectification term. Takes on different values depending on the worm's locomotion direction (whether the worm is moving forwards or in reverse). |
| $c_{vT}$ | Locomotion direction rectification parameter. Determines how much the neural activity depends on the animal's locomotion direction. |
| $c_v$ | Velocity parameter |
| $c_{\theta h}$ | Head curvature parameter |
| $c_p$ | Feeding parameter |
| $s$ | Exponentially weighted moving average (EWMA) timescale parameter. |
| $b$ | Baseline activity parameter |
| $n[0]$ | Initial condition parameter |
| $\sigma_{noise}$ | White noise parameter |
| $\sigma_{SE}$ | Autocorrelative noise parameter |
| $\ell$ | Autocorrelative noise timescale parameter |
| $\mathcal{GP}$ | Gaussian process |
| $K_{GN}, K_{SE}$ | Gaussian process kernels |
| $\tau_{1/2}$ | Half-decay parameter that explains how far back in time each neuron integrates behavior. It is not directly included in the model, but is derivable from the EWMA timescale parameter $s$ and is reported below due to its more intuitive meaning. |

We fit this model (The *C. elegans* Probabilistic Neural Encoding Model, or CePNEM) on all neurons in all recordings and found significant encoding of at least one behavioral feature in 1,168 total neurons (83 ± 10 out of an average of 143 total neurons recorded per animal; 14 animals total; example neurons shown in Fig. 1N; additional examples with cross validation shown in Fig. S2C; see Methods for description of statistics). We performed several control analyses to ensure that these results reflected genuine behavioral encoding, rather than motion or model fitting artifacts. First, we performed the same procedure on animals expressing pan-neuronal GFP rather than pan-neuronal GCaMP. Using the same statistical criteria applied above, only 2.1% of neurons significantly encoded behavior in any of these GFP animals (versus 58.6% in GCaMP datasets; Fig. S2D). We were also concerned that the model could potentially explain neural activity via overfitting, despite our efforts to calibrate the noise model (see Methods). However, we found that fitting neural activity from one animal using the behavioral features from other animals (i.e. a scrambled control) resulted in only 2.7% of neurons encoding this incorrect behavior, suggesting that the model was unable to use overfitting to explain neural activity (Fig. S2D). This is consistent with our finding that the model shows a high level of cross-validated performance, exceeding simpler versions of the model with fewer predictor terms (Fig. S2A, Fig. S2C).

Finally, we examined the extent to which these model fits captured the overall variance in neural activity across the brain that was related to overt behavior. There were indeed neurons with evident calcium dynamics not well fit by CePNEM, but it was ambiguous whether these neurons encoded behavior in a manner not captured by CePNEM or, alternatively, whether their activity was related to other ongoing sensory or internal variables. To distinguish between these possibilities, we examined the model residuals, which are the neural activity across the brain unexplained by CePNEM. As a generic test to see whether these residual dynamics reflected behavior, we attempted to decode behavioral features using all neurons’ model residuals and, as a control, we also attempted to decode the same features using the original neural activity traces. As is shown in Fig. S2E, decoding from the full neural traces was highly successful, but decoding from the model residuals was close to chance levels. This was also true for behaviors not explicitly included in the model (angular velocity and body curvature), suggesting that the model was able to construct information about neural encoding of those behaviors out of the behaviors in the model (velocity, head curvature, and feeding). For example, angular velocity could be constructed by encoding head curvature over a longer timescale. While we cannot rule out that these residuals are related to a motor program that we were unaware of, these data suggest that neural variance unexplained by the model is largely unable to predict behavior. Therefore, the model captures the majority of the neural variance related to overt behavior.

### Diverse representations of behavior across the *C. elegans* brain

We used the results of our CePNEM fits to perform a global characterization of how the full set of neurons we recorded across each animal’s brain encode its behavior. Among the recorded neurons, encoding of velocity was most prevalent, followed by head curvature and feeding (Fig. 2A). 58% of the recorded neurons encoded at least one behavior (Fig. 2B), with approximately one third of these conjunctively encoding multiple behaviors (Fig. 2B). The neurons varied in how much they weighed current versus past behavior. The majority of the neurons primarily encoded current behavior, but a sizeable subset strongly weighed past behavior (Fig. 2C; 20% of the neurons had an exponential half-decay of *τ*_1/2_ > 12.5sec, much longer than GCaMP7f sensor half-decay of <1 sec). Long timescale encoding was most prominent among forward-active velocity encoding neurons (Fig. S2G). This suggested that current neural activity may contain information about past velocity. Indeed, we found that we were able to train a linear decoder to predict prior velocity (up to at least 20 sec prior) based on current neural activity (Fig. 2D; this was not due to current velocity simply predicting past velocity, see black line in Fig. 2D). These data indicate that neurons can exhibit many different representations of past and present behavior, captured by the CePNEM model.

Using our full set of recordings and model fits, we first analyzed how velocity was represented across the full set of neurons. As described above, neurons that encode a given behavior can be tuned to that behavioral feature in a variety of ways. For example, a neuron that represents velocity could encode analog velocity, binary movement direction, and more. Using the statistical framework afforded by our fitting procedure, we determined whether each velocity-encoding neuron carried analog information about velocity during reverse and forward movement and, if so, whether the slope of its tuning was positive or negative. These slopes were computed from each possible set of model parameters from that neuron’s CePNEM fit, each of which corresponds to a particular slope value; statistical tests then checked whether the null hypothesis of zero slope could be rejected. Combining all possible reverse and forward tunings, there were eight ways that a neuron could be tuned to velocity (Fig. 2E). Most neurons (87%) exhibited rectified tunings, in which the encoding of forward and reverse speed differed. A smaller set of neurons represented velocity in a fully analog fashion and, as described above, others encoded slow locomotion. To highlight how our model fits accurately capture the dynamics of neurons with different tunings, Fig. 2H shows five different simultaneously recorded neurons that all showed higher activity during forward movement, yet their dynamics during forward and reverse behaviors are quite different (note dynamics, rise times, etc). Fig. 2H shows the CePNEM fits to each neuron, revealing how they encode velocity with different tunings and timescales. Altogether, these results show that representations of velocity are diverse among the neurons, and that CePNEM can accurately describe each of these types of representations.

We were also able to accurately model the neurons that encode head curvature, which controls the steering of the animal during navigation (Fig. 2F). We found that across the full set of recorded neurons 90% of the neurons that encoded head curvature did so in a manner that depended on movement direction (i.e. these neurons had significant encoding of both head curvature and movement direction). Thus, we categorized this full set of neurons based on both their head curvature tuning (activity during dorsal versus ventral head bending) and velocity tuning (activity during forward versus reverse movement). Most neurons only displayed head curvature-associated activity changes during forward or reverse movement, with more neurons in the forward-rectified group (Fig. 2F; see examples in Fig. 2I). Ventral-active neurons were slightly more prevalent than dorsal-active neurons, which could be related to the ventral bias of omega bending by *C. elegans* (Croll, 1975). This indicates that the network that controls head steering in *C. elegans* is broadly impacted by the animal’s movement direction, which could relate to the asymmetry in how animals must steer during forward versus reverse movement. Note that because we examined tuning to the same range of head angles during forward and reverse movement, these differences are not simply explained by a different range of head angles being explored during forward versus reverse movement. In addition to these neurons that encode the animal’s acute head curvature, a smaller group of neurons encoded angular velocity (head curvature integrated with a half-decay parameter >5sec), sometimes in combination with the animal’s movement direction (Fig. S2F shows examples).

Neural representations of the animal’s feeding (i.e. pharyngeal pumping) rates were also diverse (Fig. 2G; examples in Fig. 2J). Many neurons displayed analog tuning to feeding rates. In addition, a separate set of neurons encoded feeding in conjunction with movement direction, such that their tuning to feeding differed depending on movement direction. Neurons could be positively or negatively correlated with feeding.

The above analyses suggest a surprising amount of heterogeneity in how *C. elegans* neurons encode behavior. Neurons that encode single behaviors like velocity have a wider range of tunings and timescales than was previously known, and there is more extensive combinatorial encoding than expected also. To obtain a more complete and continuous view of these different representations, we used UMAP to embed the neurons into a low-dimensional subspace, where the proximity between the neurons indicates how similarly they encoded behavior (all neurons from all animals in Fig. 3A; data from single animals embedded in the same UMAP space in Fig. S3A-B; GFP controls in Fig. S3C; median CePNEM fits only (i.e. one dot per neuron) in Fig. S3D; see Methods for details). This analysis could in principle reveal distinct clusters of cells, which would correspond to discrete subgroups of neurons that encode behavior the same way. Alternatively, the neurons could be evenly distributed in the subspace without any clusters if the representations were more heterogeneous and varied. As is shown in Fig. 3A, the neurons were diffusely distributed in the subspace, with no evident clustering. Examining where neurons with different encoding types were localized in this subspace revealed a basic organization of how neurons were arranged (Fig. 3B-E). Encoding of forward versus reverse velocity was graded along one axis, and encoding of feeding was graded along the other. Head curvature and timescale information were more distributed. The continuous, rather than clustered, nature of the distribution of neurons was especially evident when examining locations of neurons with different tuning curves. For example, neurons with different tunings to forward velocity were represented along a continuum in one region of the plot (Fig. 3F). However, UMAP projections can be sensitive to parameters, so we also examined whether neurons were clustered versus continuous using standard metrics for data clusterability. Indeed, these analyses suggested that the neurons that were not clusterable into discrete groups (Fig. S4A; optimal number of clusters was two, the minimum allowed by the metric). These results suggest that the boundaries between neurons in terms of their representation of behavior are mostly continuous rather than discrete. Thus, rather than sorting into discrete modules, the neurons represent behavior along a continuum.

**Figure 3.**
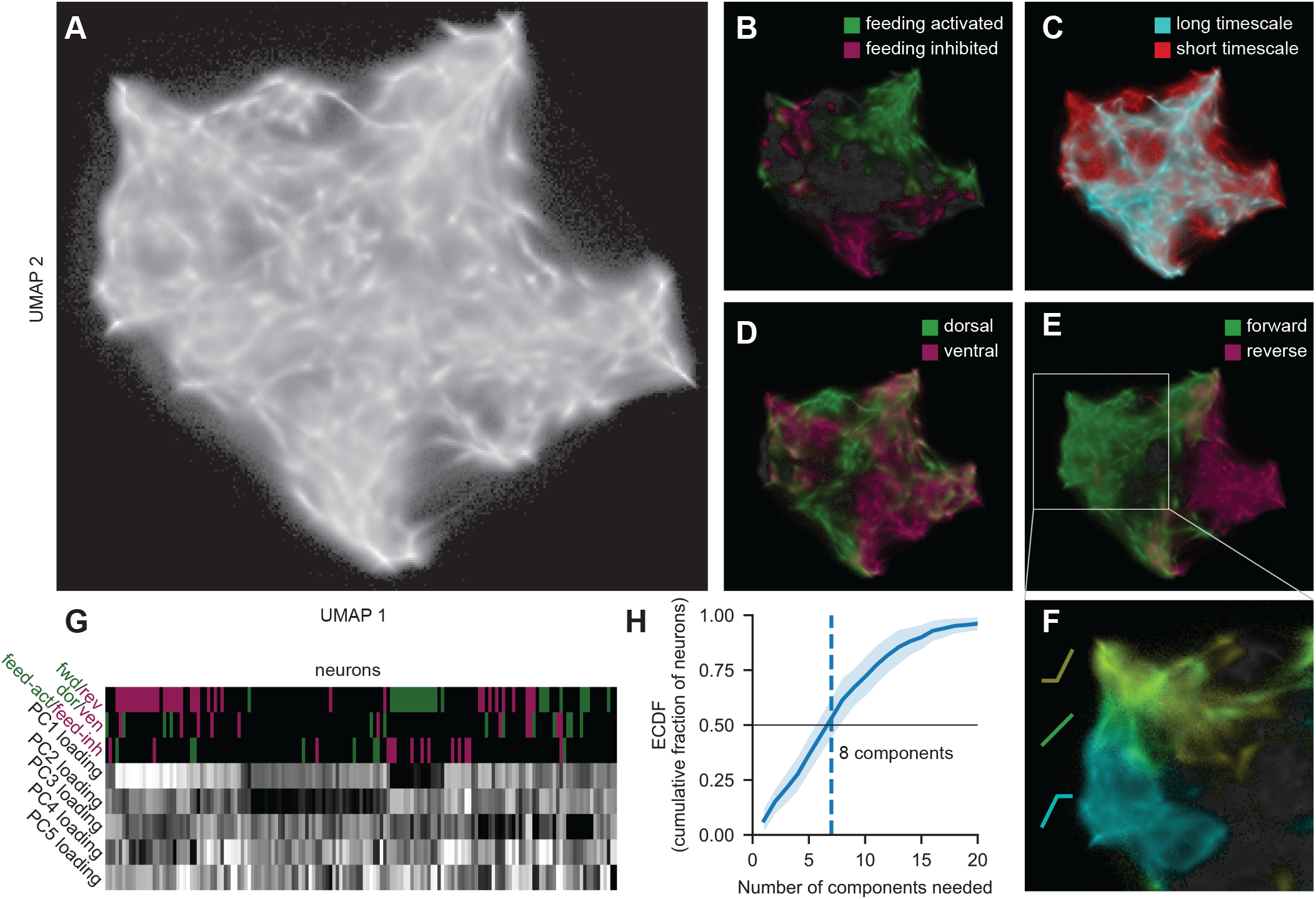
Global analysis of the encoding properties of neurons across the *C. elegans* nervous system. (A) Low-dimensional UMAP embedding space for neurons. Neurons were embedded in this space using a similarity metric that computes how similarly the neurons encode behavior; thus, proximity indicates encoding similarity (see Methods for details). In this plot, we projected all points from all CePNEM posteriors in 14 animals into this defined UMAP space to create the point cloud here, which is shown using a log scale for brightness. Fig. S3D shows just the median fits from the posteriors (i.e. one dot per neuron) projected into this space, which yields the same tiling and shape. See Methods for more details. (B-E) UMAP space where neurons are color coded by their behavioral encodings: feeding activated vs inhibited (B), long (>20 sec) vs short (<20 sec) half-decay times (C), dorsal vs ventral (D), and forward vs reverse (E). (F) Enlarged view of the portion of the UMAP space containing forward neurons, where the neurons are color-coded by the type of forward velocity tuning they have (tunings are depicted next to neuron groups in corresponding colors). (G) Neurons from an example animal, together with their tuning to velocity, head curvature, and feeding, and loadings onto the top five principal components in an example animal. The neurons are hierarchically clustered by their loadings onto the PCs, which causes neurons with similar tunings to behavior to end up clustered together due to behavioral information present in the PCs. Note also that each neuron has strong factor loadings from multiple PCs (only five of which are shown here). (H) Number of principal components needed to explain 75% of the variance in a given neuron, averaged across neurons in 14 animals. Only neurons with sufficiently high signal (see Methods) are shown, to prevent this curve from being right-shifted due to explaining measurement noise. Half of the neurons require >8 principal components to explain 75% of their variance. Data are shown as means and standard deviation across animals.

How do these diverse representations of behavior arise in the nervous system? Previous work has suggested that activity in the *C. elegans* nervous system can be decomposed into different modes of dynamics that are shared by neurons (Kato et al., 2015), identifiable through Principal Component Analysis (PCA). Performing PCA on our neural datasets revealed many distinct modes of dynamics: the first three PCs explained an average of 42% of the variance in neural activity and an average of 18 PCs were required to explain 75% of the variance in neural activity (Fig. S4B). While individual PCs were related to behavioral variables, there was not necessarily a clear mapping of each PC onto single behavioral variables. Neurons can be described as linear combinations of these PCs. The relative weighting of each PC by a given neuron reflects how much it expresses each of the main modes of dynamics. In principle, a given neuron could express a single mode of dynamics or complex mixtures. We found that the neurons were almost exclusively complex mixtures of many modes of dynamics (Fig. 3G-H). Moreover, the weights of the PCs on different neurons were diverse; hierarchical clustering of these data revealed very little structure, further suggesting that there were not clear subgroups of neurons with divisions between them. However, as expected, the factor loadings were still predictive of the encoding type of the neurons, suggesting that the form of behavior representation by a neuron is constructed by how it weighs the different shared modes of dynamics in the nervous system (Fig. 3G). We note that despite this complexity, these modes of dynamics and representations of behavior could still be highly stereotyped for individual neurons classes (see below). Overall, these results suggest that there are many ongoing modes of dynamics shared among neurons. The form of behavior representation by these neurons reflects which of these modes of dynamics each neuron expresses. Because each neuron carries a complex mixture of dynamics, the neurons carry distinct representations of behavior.

### An atlas of how the defined neuron types in the *C. elegans* connectome encode behavior

We next sought to map these diverse representations of behavior onto the defined cell types of the *C. elegans* connectome. Thus, we collected additional datasets in which we could determine the identity of each recorded neuron. In these experiments we utilized the previously described NeuroPAL transgene (Yemini et al., 2021). NeuroPAL animals express three different fluorescent proteins (NLS-BFP, NLS-OFP, and NLS-mNeptune) under well-defined genetic drivers, which makes it easy to determine neural identity based on neuron position and multispectral fluorescence. They also express pan-neuronal NLS-TagRFP-T, but have no green fluorescence. We crossed the pan-neuronal NLS-GCaMP7f transgene to the NeuroPAL transgene (we used *otIs670*, a low brightness integrant of NeuroPAL shown to be phenotypically wild-type in many respects). Data were collected under the same conditions used above, except at the end of each freely-moving GCaMP recording animals were immobilized by cooling. We then collected NeuroPAL data in each fluorescent channel and registered the immobilized images back to the freely-moving images (example image in Fig. S5A).

We collected data from 12 freely-moving NeuroPAL/GCaMP7f animals. Behavior encoding was qualitatively similar in this strain, compared to the datasets described above: a similar number of neurons encoded behavior (49.5%, compared to 58.6% above), and the projections of neurons into UMAP space based on behavior encoding yielded indistinguishable results (Fig. S3B). Using NeuroPAL labels, we determined the identities of 96 ± 14 recorded neurons per animal. In total, we recorded data from 77 of the 80 neuron classes in the head. While most neuron classes are a single left/right pair of neurons, 13 of these classes consist of two or three pairs of neurons positioned in 4- or 6-fold symmetric arrangements. In such cases, we separately analyzed each of the neuron pairs, since their functions could differ (left/right pairs were pooled for all neuron classes). Thus, in total we separately analyzed the functional properties of 91 different neuron types, with an average of 11.1 neurons recorded per type. We generated CePNEM fits to all of these recorded neurons to reveal how each neuron class encodes behavior (Fig. 4A; Fig. S4B provides further explanation of how to read the atlas; Fig. S4H shows locations of many neurons in UMAP space). For neurons with previously defined encodings, our results provided a clear match to previous work: AVB, RIB, AIY, and RID encoded forward movement; AVA, RIM, and AIB encoded reverse movement; and SMDD and SMDV encoded dorsal and ventral head curvature, respectively (Gordus et al., 2015; Kaplan et al., 2020; Kato et al., 2015; Li et al., 2014; Lim et al., 2016; Luo et al., 2014; Roberts et al., 2016).

**Figure 4.**
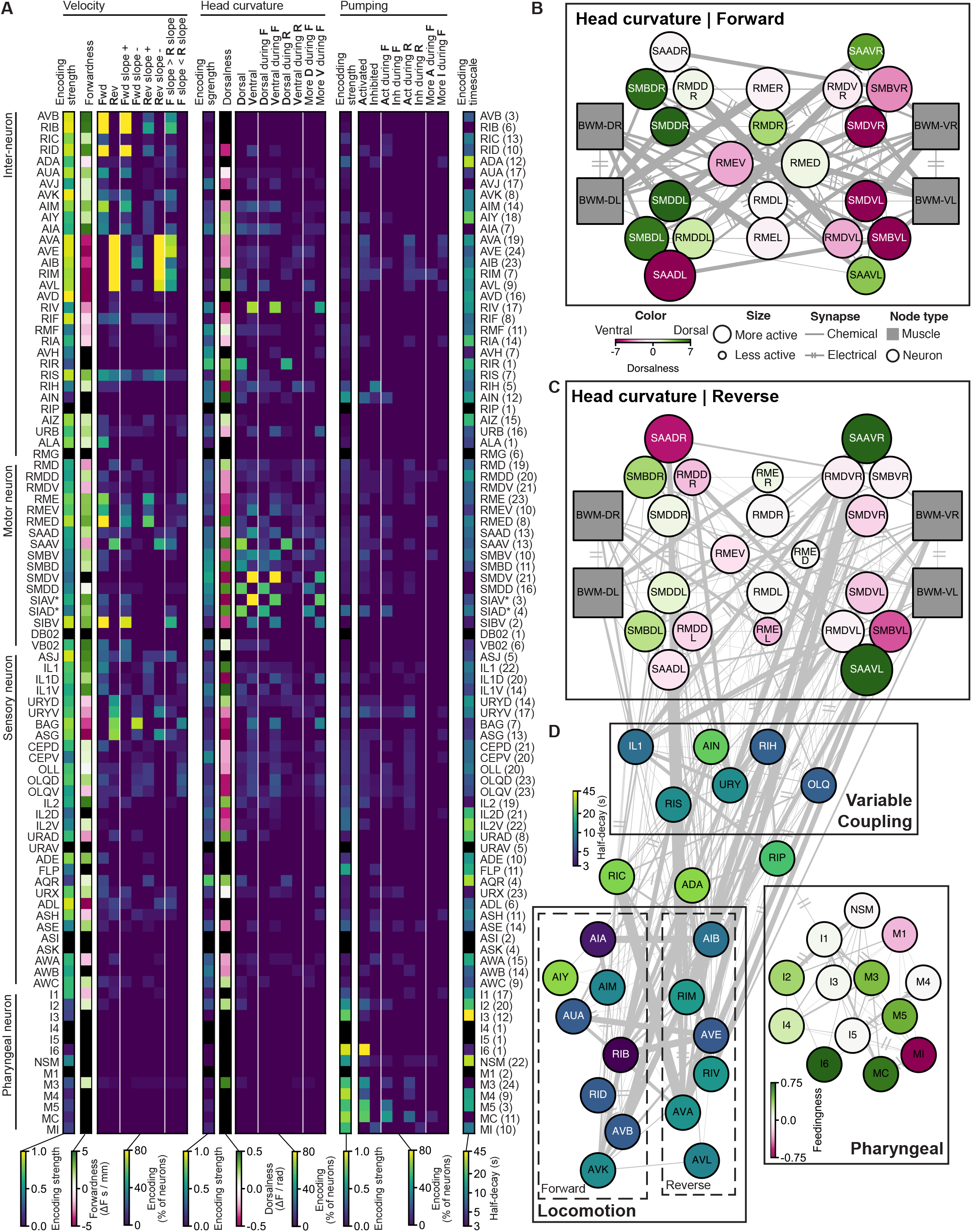
An atlas of how the different *C. elegans* neuron classes encode behavior. (A) An atlas of how the indicated neuron classes encode features of the animal’s behavior, derived from analysis of fit CePNEM models. All neuron classes that were recorded, identified, and mapped back to GCaMP traces are shown. Columns indicate the encoding features of the neurons, as follows (see Methods for additional details):

- **Encoding strength** for each of the three behaviors indicates an approximation of the relative variance in neural activity explained by each behavioral variable.
- **Forwardness** and **Dorsalness** capture the steepness of the tuning to velocity and head curvature, respectively.
- **Encoding timescale** is the median exponentially weighted moving average (EWMA) half-decay time from the CePNEM model, which indicates how the neuron’s activity weighs past versus present behavior for the parameter(s) that it encodes. All other columns show the fraction of recorded neurons that significantly encoded behavior defined by each column description (see Methods for more details, and see Figure S5B for a legend that explains how these encodings relate to the neural tunings from Figure 2):

- **Fwd**, **Rev**, **Dorsal**, **Ventral**, **Activated**, and **Inhibited** represent neurons with that overall tuning to the behavior (velocity, head curvature, or feeding).
- **Fwd slope** -, **Fwd slope +**, **Rev slope** -, and **Rev slope +** represent neurons with that slope in their tuning curves to velocity during the specified movement direction. For example, a neuron with both **Fwd slope** - and **Rev slope +** would have a tuning curve to velocity that looks like Λ, ie encoding slow locomotion.
- **F slope > R slope** and **F slope < R slope** represent neurons displaying rectification in their velocity tuning curves, with the slope of the tuning curve during forward movement being either larger or smaller than during reverse movement, respectively.
- **Dorsal during F**, **Ventral during F**, **Dorsal during R**, **Ventral during R**, **Act during F**, **Inh during F**, **Act during R**, and **Inh during R** represent neurons with the specified tuning to the behavior (head curvature or feeding) during the specified movement direction (**F**orward or **R**everse).
- **More D during F**, **More V during F**, **More A during F**, and **More I during F** represent neurons with different tunings to the behavior (head curvature or feeding) during forward versus reverse locomotion. For instance, a **More D**(orsal) **during F**(orward) neuron would indicate a neuron with stronger dorsal-tuning during forward movement compared to reverse; the other categories behave similarly. Black pixels mean that the neuron was never found to significantly encode the behavioral parameter. Numbers in parenthesis on the right indicate the number of CePNEM fits for that neuron class (left/right neurons are counted separately; first and second halves of videos, which have different model fits, are also counted separately). (B-C) Circuit diagram of the neuron classes that innervate the dorsal and ventral head muscles, with colors (tuning of neuron to dorsal versus ventral head curvature) and circle sizes (overall activity level during forward or reverse) indicating how each neuron is tuned to behavior during forward (B) and reverse (C) movement. Grey connections are from the *C. elegans* wiring diagram. Note the large degree of symmetry in neural encoding and the shift in tuning/activity based on forward (B) versus reverse (C) movement. (D) Circuit diagram of several sub-circuits including locomotion (forward/reverse) and pharyngeal circuit, connected to the head curvature circuit in (C). The color of each node indicates the median half-decay time of the neuron class. The color in the pharyngeal circuit indicates the feedingness (steepness of neuron’s tuning curve to feeding). The middle group (AIN, RIH, RIS, URY, IL1, OLQ) are richly connected to locomotion, head curvature, and feeding (via the RIP neuron that links to the pharyngeal circuit).

This analysis revealed many new features of how the *C. elegans* nervous system is organized to control behavior. Although the velocity circuit has been fairly well studied, many new features still emerged (Fig. 4A, 4D). The neurons that encode forward movement displayed a wider range of rectified and non-rectified representations of velocity than was previously known, and included many neurons not previously implicated (AIM, AUA, and others). The reverse neurons were more uniform in their tunings to velocity, but several of them also represented head curvature during reverse movement (AVL, RIV), suggesting that they may control aspects of bending/turning during reverse movement. Neural representations of velocity also spanned multiple timescales. RIC, ADA, AVK, AIM, and AIY represented velocity over long timescales, showing encoding of the animal’s recent velocity over tens of seconds (half-decays of 10-30s). RIC and ADA form dense synaptic outputs onto the command neurons that drive the acute forward/reverse movement of the animal (Chalfie et al., 1985; Gordus et al., 2015; Kato et al., 2015; Roberts et al., 2016), suggesting that they may integrate recent behavior and influence current behavior. We silenced some of these neurons newly implicated in velocity control (AIM, RIC) and found that this indeed altered animals’ velocity, but it did not perturb head curvature or feeding behaviors, consistent with the notion that the neurons are involved in velocity control (Fig. S5C).

These data also revealed for the first time how neural activity is coordinated in the circuit that controls head steering during navigation. The neuron classes in this network are often 4-fold symmetric, consisting of separate neuron pairs that innervate the ventral and dorsal head muscles to allow for steering. These opposing dorsal and ventral neuron classes were identified as being functionally antagonistic in our analysis (Fig. 4A-C). Strikingly, our analysis showed that the neural control of head steering is dramatically different during forward versus reverse motion (Fig. 4B vs Fig. 4C). Some of the neuron classes that encode head curvature are selectively active during forward (RMED/V) or reverse (SAAV) movement. Other classes have more robust tuning to head curvature during forward movement (SMDD/V, SMBD/V). In addition, RMDD was more active during dorsal head bending during forward movement, but switched to prefer ventral head bending during reverse movement. The forward-rectified tuning of the SMD neuron class has been previously described and matches our results (Kaplan et al., 2020). Our data here now show that this entire network shifts its functional properties depending on movement direction. This suggests that the network likely performs different sensorimotor transformations as animal steer towards a target during forward movement compared to when they move away from a target during reverse movement. Our data also show that the timescale of representation of head curvature differs among the neurons. SMD neurons encoded head bending on a rapid timescale such that their activity faithfully tracked head oscillations. However, the other classes (RME, RMD, SMB, and SAA) had longer timescale integration, such that their activity both represented acute head oscillations and longer timescale changes in dorsal/ventral bending (i.e. gradual steering signals).

We also identified other functional groups with novel features (Fig. 4D). For example, most of the neurons that encoded feeding were in the pharyngeal nervous system, but several extrapharyngeal neurons also contained information about feeding, such as RIH, AIN, and SIA. We also found that many neurons that had not been well studied in the literature had variable tunings to different motor programs that either differed across animals and/or were highly multiplexed in individual animals. These neurons (AIN, OLQ, IL1, RIH, URY, others) appear to be able to flexibly couple to different motor circuits in the animal. We confirmed that our NeuroPAL labeling procedure and our registration methods for these neurons were determined with equal confidence to the other neuron classes, suggesting that identification errors are unlikely to explain these observations (Fig. S5D,E; Fig. S5G shows example data). Further supporting this, these neurons also changed encoding over the course of individual continuous recordings (see below). These results thus identify many neuron classes in the *C. elegans* nervous system that can flexibly couple to different behavioral circuits. Overall, these datasets have now provided a functional map of how most neuron classes in the *C. elegans* nervous system encode the animal’s behavior.

### The encoding of behavior is dynamic in many neurons, and is influenced by the behavioral state of the animal

While examining these datasets, we noted that in several cases the encoding properties of neurons appeared to change over time in a single recording. Therefore, we systematically analyzed our data to determine whether neural representations of behavior dynamically change. To accomplish this, we fit two CePNEM models trained on the first and second halves of the same neural trace and used the Gen statistical framework to assess whether the model parameters had significantly changed between these time segments. In addition, a neuron’s encoding was only considered to have changed if another model trained on the full time range performed significantly worse on the first and seconds halves of the data, compared to the models trained on these halves (see Methods). To ensure that model overfitting would not result in the spurious detection of encoding changes, we ran this analysis on simulated neurons from our model with constant ground-truth parameters and verified that our statistical approach did not detect any changes in encoding in these simulated neurons (Fig. S6A; photobleaching was also ruled out as a contributing factor, see Fig. S6B and Methods).

We observed that ~20% of neurons that encoded behavior changed that encoding over the course of our continuous neural recordings. Some examples of these neurons are shown in Figure 5B. This suggests that a sizable portion of the *C. elegans* nervous system is flexible, with many neurons changing how they map onto behavior over time, even in a 16-minute recording. We found a similar fraction (14%) of neurons change encoding in the NeuroPAL strain, so we determined the ground-truth identities of these flexible neurons. This analysis revealed a set of neurons that significantly overlap with those that variably encode behavior across animals (Fig. 5G, neurons on right; compare to ‘variable coupling’ neurons in Fig. 4D). This suggests that these flexible neurons can couple to different circuits and change how strongly they couple to those circuits over time. Motor neurons were unlikely to change their encoding, while sensory neurons and interneurons with high amounts of sensory input were more likely to change encoding (Fig. 5G). Moreover, specific neuron classes appeared to show different magnitudes of encoding changes: OLQD’s encoding changed drastically in some cases (Fig. 5D; Fig. 5E shows how OLQD’s encoding moved through UMAP encoding space overall, and in individual recordings), while AVE only showed low magnitude encoding changes (Fig. 5D-E) and always remained tuned to reversal. Among all the neurons, many different types of encoding changes were observed: changes in which behaviors were encoded by a neuron, including complete losses of behavior encoding and swaps in which behaviors were encoded; and more subtle changes in tuning to the same behavior (Fig. 5F). Overall, these results suggest that a defined subset of neurons in the higher layers of the *C. elegans* connectome can be variably coupled to behavioral circuits and remap how they couple to these circuits over time.

**Figure 5.**
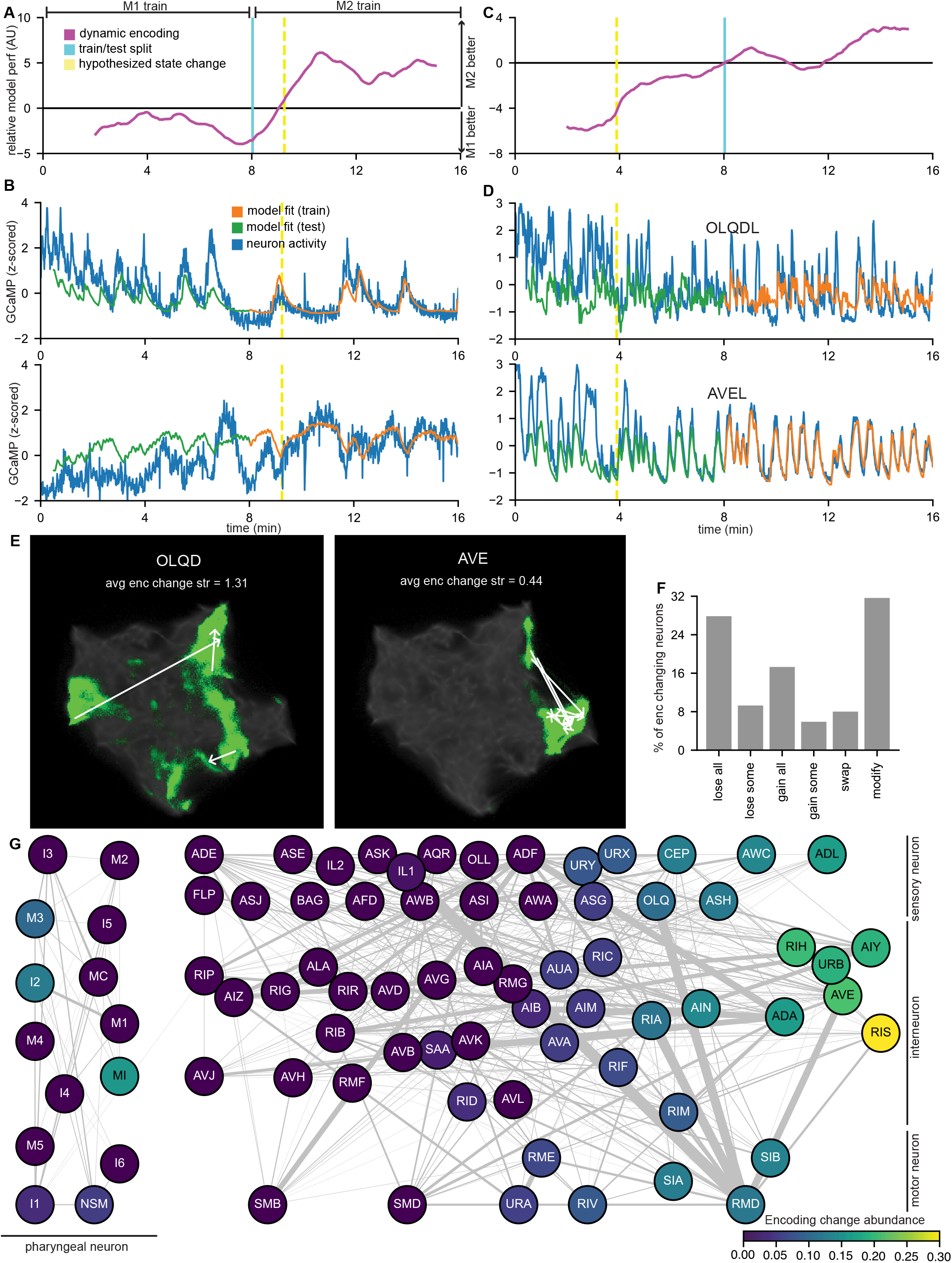
Neural representations of behavior dynamically change over time at stereotyped sites in the *C. elegans* connectome. (A) Data from an example animal showing a sharp change in relative model performance in flexible encoding neurons. The relative model performance (y-axis) was calculated as follows. We fit two CePNEM models (M1 and M2) to the first and second halves of the 16min recording, respectively. We then computed the difference between the errors of the two median model fits (computed as the MSE to the observed neural trace) at every time point, and took a moving average over 200 time points. This was then averaged across neurons. As is indicated, values >0 indicate that M2 model fits better, whereas values <0 indicate that M1 model fits better. A sudden change in this metric (dashed yellow line) indicates a sudden shift in which model fits the neural data, indicative of an encoding change. (B) Two example neurons from the animal in (A) with CePNEM fits, showing a change in neural encoding of behavior at the moment of the hypothesized state change. The models were fit on the parts of the data where the model fit is colored in orange and used to predict activity over the full time series. (C) Data from an example NeuroPAL animal that also shows a sharp change in relative model performance, displayed as in (A). (D) Example neurons OLQDL and AVEL with median CePNEM fits from the animal in (C). (E) UMAP plots to show the degree to which neurons change encoding. Each plot shows the projections of all CePNEM fit posteriors across all recordings for OLQD (top) and AVE (bottom) into the UMAP space, in green. The background UMAP space of all neurons is shown in grey. White arrows are drawn between the median model fits between two time ranges from individual recordings if the neuron had an encoding change between those time ranges. Average encoding change strength (indicated) was computed as the variance of the difference between the two time-range specific median CePNEM fits extrapolated over the full time range, divided by the maximum of the variances of those two extrapolated fits, averaged over all animals where the neuron in question had an encoding change. See Methods for more details. (F) A categorization of how the encoding properties of neurons can change across all recorded SWF415 animals. More detailed explanations of each category, which were computed based on comparing which behaviors the neuron encoded before and after the encoding change: “lose all” (the neuron lost all of its tuning to behavior), “lose some” (the neuron lost tuning to at least one behavior, and didn’t gain tuning to any behavior), “gain all” (the neuron did not encode any behavior before the encoding change, but did afterwards), “gain some” (the neuron gained tuning to at least one behavior, and didn’t lose tuning to any behavior), “swap” (the neuron both gained and lost tuning to behaviors, effectively switching which behavior it encoded), and “modify” (the neuron encoded the same set of behaviors, but in a different way). (G) A diagram of all neurons detected in our NeuroPAL recordings, sorted by neuron type and encoding change abundance. Encoding change abundance was computed as the average encoding change strength as computed in (E) over all datasets where that neuron was detected. Interneurons are vertically sorted by the fraction of their inputs that come from sensory neurons.

We next sought to understand the temporal structure of these encoding changes. For instance, individual neurons could remap independently of each other, or there could be a circuit-wide, temporally-synchronous shift. To address this, we developed a metric to identify when an encoding change took place. This was computed by subtracting the errors of models trained on different time regions of the same neural trace, and averaging this metric across all neurons that encode behavior in that animal (Fig. 5A and 5C, purple line; additional controls in Fig. S6C and S6D; see Methods). Sharp changes in this metric should reveal time points where the relative performance of the models change. If neurons change encoding independently, averaging across neurons will smooth out any individual encoding changes, and the metric will gradually increase. However, if there is a synchronous encoding shift across the brain, the metric will suddenly change at the time of that shift. Intriguingly, while we did observe instances of non-synchronized encoding changes (see Fig. S6E), we also observed that in many cases there was a synchronous change across many neurons (Fig. 5A, Fig. 5C). By examining our NeuroPAL datasets, we found that certain neuron classes were more likely to change encoding at the same time as one another (Fig. S6K). In addition, the number of neurons that changed encoding was positively correlated with the degree of behavioral change across the hypothesized moment of the change (Fig. S6L). Overall, these results suggest that at times there is a coordinated remapping where many neurons change how they represent behavior.

These synchronous shifts in the neural encoding of behavior might reflect ongoing changes in the animal’s internal or behavioral state. Alternatively, they might be explained by a sudden change in the animal’s sensory surroundings, despite our efforts to record animals in arenas with homogeneous sensory cues. We performed additional experiments to directly test whether changing the animal’s behavioral state could elicit a synchronous encoding change across neurons. Behavioral states are typically defined as persistent changes in behavior that outlast the sensory stimuli that initiate them (Anderson and Adolphs, 2014; Flavell et al., 2022). Previous work has shown that aversive stimuli can induce this type of behavioral change in *C. elegans* (Ardiel et al., 2017; Byrne Rodgers and Ryu, 2020; Chew et al., 2018). Therefore, we delivered a sudden, noxious heat stimulus to animals part way through our recordings (Figure 6A,E). Specifically, we recorded 11 additional datasets in which we used a 1436-nm laser to heat the agar around the worm’s head by 10°C for 1 sec after 4-6 minutes of baseline recording (Fig. 6A; temperature decayed back to baseline with a time constant of 0.39s, fully returning to baseline within 3 sec). Heat stimuli of this amplitude are known to activate the AFD, FLP, and AWC sensory neurons in immobilized animals (Kotera et al., 2016). We found that this brief, aversive stimulus elicited an immediate avoidance (reversal) behavior and a sharp reduction in feeding (Fig. 6B). Animals continued to exhibit reduced feeding rates and increased reversal rates for many minutes after the stimulus, suggesting that the transient heat stimulus induces a persistent behavioral state change (Fig. 6B). However, animals’ behavior reverted to normal within an hour and their viability was not adversely impacted by the stimulus (Fig. 6C-D). We applied our encoding change analysis to these datasets, comparing model fits from the time period prior to the stimulus to fits from a time period after the stimulus. We found that 5% of neurons radically changed their encoding of behavior at the moment of the heat stimulation, which lasted for many minutes afterwards. This number being smaller than the fraction of neurons exhibiting encoding change in our non-stimulus datasets (20%) could either be due to the stimulation evoking a smaller, stimulus-specific set of neurons to change encoding, or simply due to the fact that our model fits in heat-stimulus datasets were given a smaller amount of data, decreasing our statistical power to identify encoding changes. The types of encoding changes in response to the heat stimulus were varied: some neurons lost their coupling to behavioral circuits and went nearly silent, others suddenly displayed tuning to behavior where there was none before, and yet others displayed tuning changes (Fig. 6G). This suggests that inducing a behavioral state change elicits a shift in the network that remaps the relationship between neural activity and behavior.

**Figure 6.**
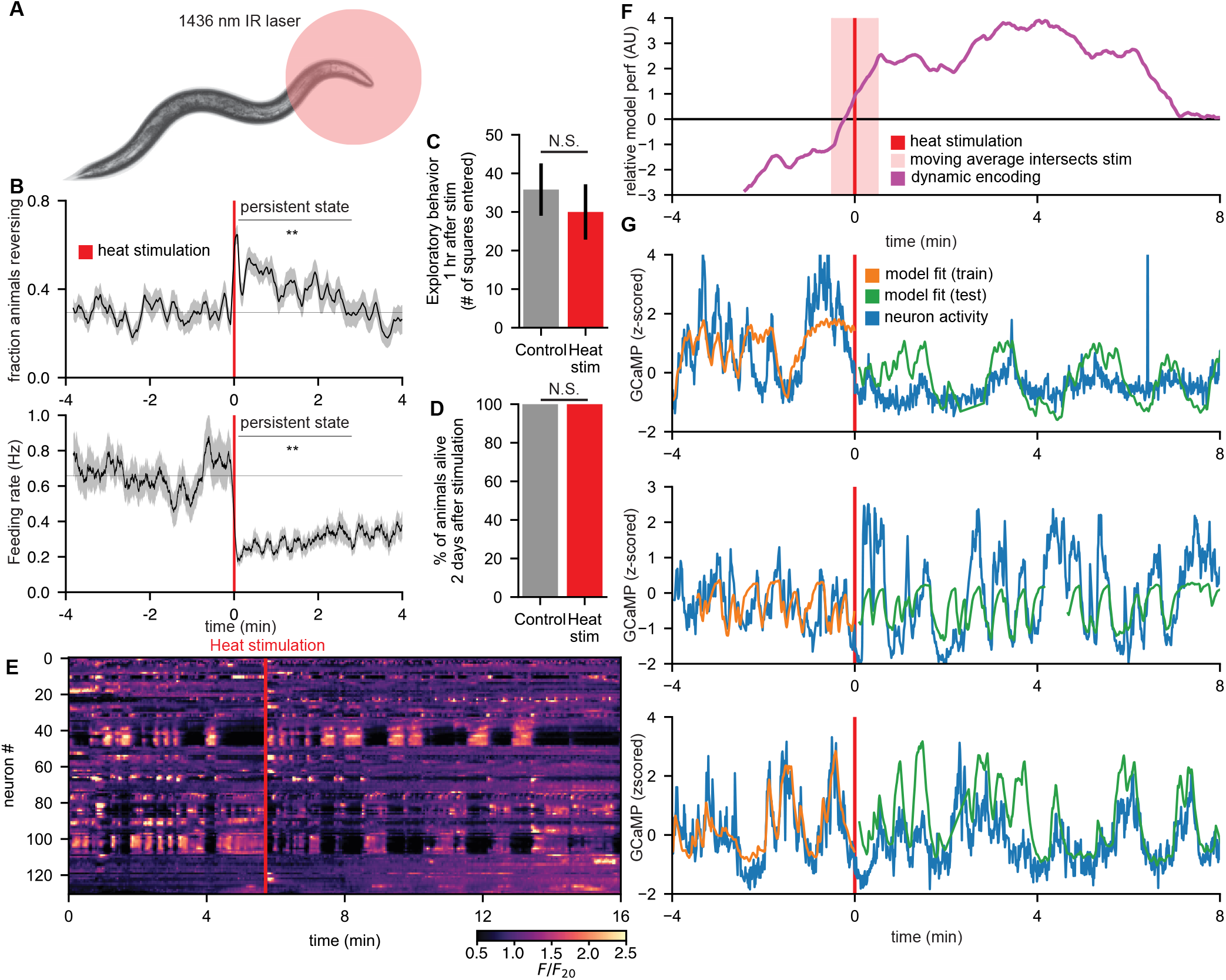
Behavioral state changes cause a widespread remapping of how neurons encode behavior. (A) A 1436nm IR laser transiently increases the temperature around the animal’s head during a whole-brain imaging session. The stimulus increased the temperature by 10°C for 1 sec and decayed back to baseline within 3 sec. This is just an illustrative cartoon. (B) Event-triggered averages of behavioral properties of 32 animals in response to the heat stimulus, demonstrating an increased reversal rate and decreased feeding rate that persist for several minutes after stimulation. **p<0.05, Wilcoxon signed rank test, comparing baseline prestimulus behavior to post-heat-stimulus behavior. (C) Experiments to examine the impact of the heat stimulation on the behavior and health of the animals. Animals subjected to the heat stimulation did not display a significant difference (***p*** = 0.62 in a Mann-Whitney U-Test computed over 10 animals) in their exploratory behavior (computed as counting the number of squares each animal entered on an assay plate) relative to mock-stimulated animals (animals that were mounted on imaging slides, but not given the thermal stimulus). (D) The heat stimulation did not kill any animals (all animals were alive 2 days after the stimulation). (E) An example 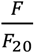 heatmap of an animal that received a heat stimulus at the indicated time (red line). Note the thermal sensory and other neurons that activate immediately after the stimulation. (F) Relative model performance (computed as in 5A, except with only 100 timepoints in the moving average) between a model trained before the heat stimulation to one trained after, demonstrating sharp change at the moment of the stimulation. Red line indicates moment of heat stimulus and shaded red region shows adjacent timepoints that could show change in relative model performance, due to a moving average sliding window that overlaps the heat stimulus. (G) Three example neurons with median CePNEM fits from animals that underwent the stimulation, showing marked and abrupt changes in their behavioral encoding immediately after the stimulus.

## DISCUSSION

Animals must adapt their behavior to a constantly changing environment. How neurons represent these behaviors and how these representations flexibly change in the context of the whole nervous system was unknown. To address this question, we developed new technologies to acquire high quality brain-wide activity and behavioral data. Using the probabilistic encoder model CePNEM, we constructed a brain-wide map of how each neuron precisely encodes behavior. By also determining the ground-truth identity of these neurons, we overlaid this map upon the physical wiring diagram. Behavioral information is richly expressed across the brain in many different forms – with distinct tunings, timescales, and levels of flexibility – that map onto the defined neuron classes of the *C. elegans* connectome.

Previous work has shown that, in both *C. elegans* and mammals, animal behaviors are accompanied by widespread changes in neural activity across the brain, resulting in a relatively low-dimensional neural space (Urai et al., 2022). This largely redundant distribution of information across the brain seems non-parsimonious. However, in this study, we found a new layer of complexity emerged when we examined the precise neural representations of behavior using high-SNR datasets and new modeling approaches. We found that while representations were complex and diverse, there was a clear logic to the encoding properties of neurons. This variability in encoding could be explained in large part by three motifs: varying timescales, non-linear (rectified) tunings to behavior, and conjunctive representations of multiple motor programs. Having many different forms of behavior representation present in the nervous system may confer this system with robustness and computational flexibility. Depending on the context, the brain may be able to selectively combine different representations to construct new coordinated behavioral outputs, dramatically expanding the effective computational space. In particular, the fact that there are diverse neural representations and that many neurons can remap their representations may allow the nervous system to generate a vast array of context-appropriate behavioral motifs. Our recording data here did not distinguish whether a given neuron’s encoding of behavior reflected the neuron causally influencing behavior or, alternatively, receiving proprioceptive or corollary discharge signals relevant to behavior; both of these types of representations are essential for a nervous system to properly control behavior.

Recent experience shapes behavior in all animals. However, how each neuron stores and uses recent historical information is poorly understood. We found that neurons can store recent motor actions with varying timescales. This allows the brain to encode the animal’s overall locomotion state at different moments in the recent past. Functionally, combining these representations with different timescales should allow the animal’s nervous system to perform computations that relate past behavior to present. We found that these representations of past behavior stretch back in time for up to a minute, but the nervous system can change over longer time scales also. In particular, we found that there is a set of neurons that can flexibly remap their relationships to behavior over many minutes. Interestingly, neurons are capable of remapping to different degrees. For example, the reversal neurons AVA and AVE are both strongly tuned to the animal’s backwards locomotion. Despite encoding the same type of information, AVA’s representation of behavior was almost completely static in our recordings whereas AVE can flexibly change its encoding. However, neurons in the sensory circuits, such as OLQ, can show even larger changes in how they encode behavior, completely switching which motor programs they encode. We found that this type of remapping occurred in a time-locked fashion in many neurons when we elicited a change in the animal’s behavioral state using a sudden aversive stimulus. This suggests that the behavioral state of the animal can remap how neurons and circuits are organized to control behavior.

Our results here reveal how neurons across the *C. elegans* nervous system encode the animal’s behavior. Even in the narrow set of environmental conditions explored in this study, we observed that ~20% of the worm’s nervous system can flexibly remap. Future studies conducted in a wider range of contexts and states will reveal whether this comprises the core flexible subset of neurons across the worm’s nervous system or, alternatively, whether the neurons that remap will be different depending on the context or state. The degree to which brain representations of behavior are constrained by synaptic wiring versus ongoing neuromodulation remains to be seen.

## ACKNOWLEDGMENTS

We thank Drew Robson, Jennifer Li, Qiang Liu, Shay Stern, Brandon Weissbourd, Nate Cermak, Robert Yang, Quilee Simeon, Cori Bargmann, and members of the Flavell lab for their comments on the manuscript. We thank Drew Robson and Jennifer Li for sharing code for spline fitting to compute posture similarity. We thank Span Spanbauer for sharing insights into model fitting techniques. V.K.M. acknowledges funding from DARPA (Award ID: 030523-00001); the Singapore DSTA/MIT SCC collaboration; and philanthropic gifts from the Aphorism Foundation and the Siegel Family Foundation. S.W.F. acknowledges funding from NIH (NS104892 and GM135413); NSF (Award #1845663); the McKnight Foundation; Alfred P. Sloan Foundation; and the JPB Foundation.

## AUTHOR CONTRIBUTIONS

Conceptualization, A.A.A., J.K. and S.W.F. Methodology, A.A.A, J.K., Z.W., E.B., M.B. D.K., J.P., V.K.M., and S.W.F. Software, A.A.A., J.K., E.B., M.B., and J.P. Formal analysis, A.A.A. and J.K. Investigation, A.A.A., J.K., Z.W., E.B., D.K., J.P., and C.E. Writing – Original Draft, A.A.A., J.K., and S.W.F. Writing – Review & Editing, A.A.A., J.K., and S.W.F. Funding Acquisition, V.K.M. and S.W.F.

## DECLARATION OF INTERESTS

The authors have no competing interests to declare.

## STAR METHODS

### Key Resources Table

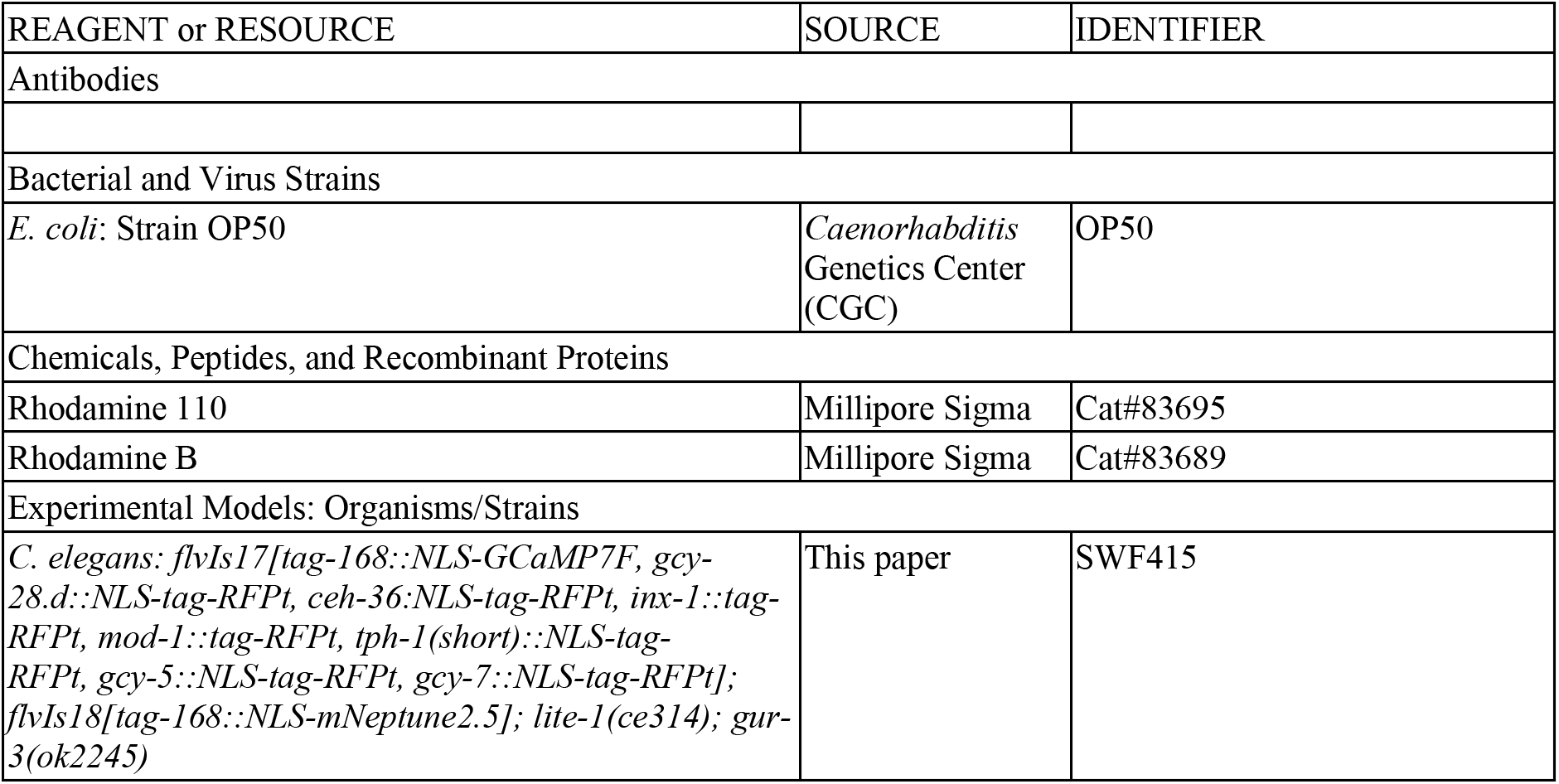
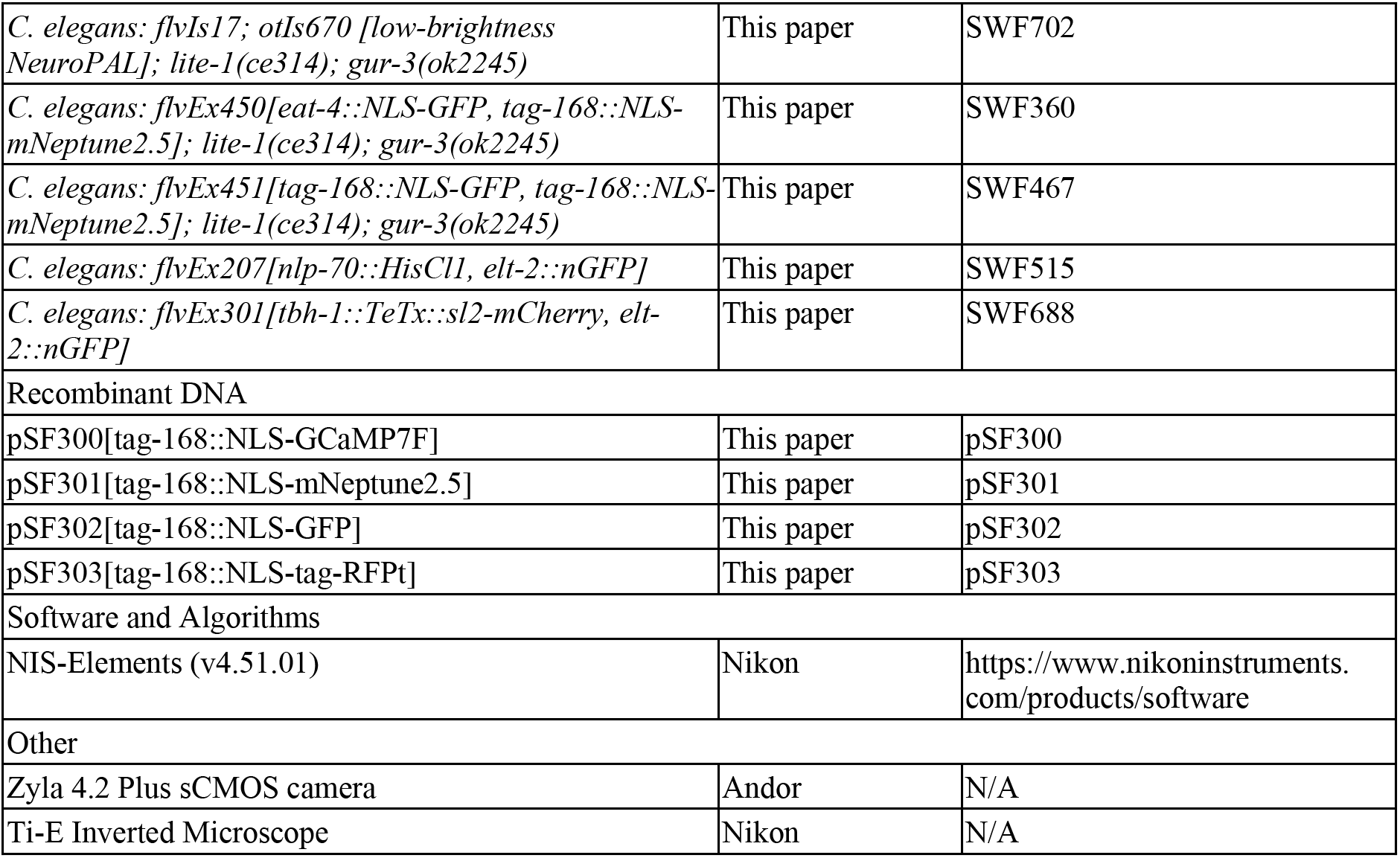

### List of key software packages used

Gen,jl, PyPlotjl, PyCalljl, HDF5.jl, ProgressMeterjl, Distributions.jl, Imagesjl, NLoptjl, DelimitedFilesjl, NaNMathjl, Clusteringjl, DataStructuresjl, Interpolationsjl, MultivariateStats.jl, Optimjl, TotalVariationjl, UMAP.jl, Lasso.jl, VideoIOjl, Imputejl, JLD2.jl, JSONjl LsqFitjl, MLBasejl, ImageTransformations.jl, HypothesisTestsjl, MultipleTestingjl, GLMjl, GLMNetjl, ForwardDiffjl, FFTW.jl, Distancesjl, DSPjl, CoordinateTransformationsjl, Combinatorics.jl, Colorsjl, ColorTypesjl, Cairojl, CUDA.jl

### Lead Contact statement

Further information and requests for resources and reagents should be directed to and will be fulfilled by the lead contact, Steven Flavell.

### Materials availability statement

All plasmids, strains, and other reagents generated in this study are freely available upon request.

### Data and code availability statement

The code used for microscope control, image processing, and data analysis is openly available at https://www.dropbox.com/s/3e5qnzam2xvdf4f/code.zip?dl=0

### Recordings of neural activity and behavior

#### Transgenic animals

Four transgenic strains were recorded in this study, as described in the text. The first (SWF415) contained two integrated transgenes: (1)*flvIs17:* tag-168::NLS-GCaMP7f, along with NLS-TagRFP-T expressed under the followed promoters: *gcy-28.d, ceh-36, inx-1, mod-1, tph-1(short), gcy-5, gcy-7;* and (2)*flvIs18:* tag-168::NLS-mNeptune2.5. The second strain we recorded (SWF702) contained two integrated transgenes: (1) *flvIs17:* described above; and (2) *otIs670:* low-brightness NeuroPAL (Yemini et al., 2021). Strains were backcrossed 5 generations after integration events. The third and fourth strains are non-integrated transgenic strains expressing NLS-GFP and NLS-mNeptune2.5 in defined neurons, listed in the Key Resources Table (SWF360 and SWF467).

#### Microscope

Animals were recorded under a dual light-path microscope that is similar though not identical to one that we have previously described (Ji et al., 2021). The light path used to image GCaMP, mNeptune, and the fluorophores in NeuroPAL at single cell resolution is an Andor spinning disk confocal system with Nikon ECLIPSE Ti microscope. Light supplied from a 150 mW 488 nm laser, 50 mW 560 nm laser, 100 mW 405 nm laser, or 140 mW 637 nm laser passes through a 5000 rpm Yokogawa CSU-X1 spinning disk unit with a Borealis upgrade (with a dual-camera configuration). A 40x water immersion objective (CFI APO LWD 40X WI 1.15 NA LAMBDA S, Nikon) with an objective piezo (P-726 PIFOC, Physik Instrumente (PI)) was used to image the volume of the worm’s head (a Newport NP0140SG objective piezo was used in a subset of the recordings). A custom quad dichroic mirror directed light emitted from the specimen to two separate sCMOS cameras (Zyla 4.2 PLUS sCMOS, Andor), which had in-line emission filters (525/50 for GCaMP/GFP, and 610 longpass for mNeptune2.5; NeuroPAL filters described below). Data was collected at 3 × 3 binning in a 322 × 210 region of interest in the center of the field of view, with 80 z planes collected at a spacing of 0.54 um. This resulted in a volume rate of 1.7 Hz (1.4 Hz for the datasets acquired with the Newport piezo).

The light path used to image behavior was in a reflected brightfield (NIR) configuration. Light supplied by an 850-nm LED (M850L3, Thorlabs) was collimated and passed through an 850/10 bandpass filter (FBH850-10, Thorlabs). Illumination light was reflected towards the sample by a half mirror and was focused on the sample through a 10x objective (CFI Plan Fluor 10x, Nikon). The image from the sample passed through the half mirror and was filtered by another 850-nm bandpass filter of the same model. The image was captured by a CMOS camera (BFS-U3-28S5M-C, FLIR).

A closed-loop tracking system was implemented in the following fashion. The NIR brightfield images were analyzed at a rate of 40 Hz to determine the location of the worm’s head. To determine this location, the image at each time point is cropped and then analyzed via a custom-trained network with transfer learning using DeepLabCut (Mathis et al., 2018) that identified the location of three key points in the worm’s head (nose, metacorpus of pharynx, and grinder of pharynx). The tracking target was determined to be halfway between the metacorpus and grinder (central location of neuronal cell bodies). Given the target location and the error, the PID controller configured in disturbance rejection sends velocity commands to the stage to cancel out the motion. This permitted stable tracking of the *C. elegans* head.

#### Mounting and recording

L4 worms were picked 18-22 hours before the imaging experiment to a new NGM agar plate seeded with OP50 to ensure that we recorded one day-old adult animals. A concentrated OP50 culture to be used in the mounting buffer for the worm was inoculated 18h before the experiment and cultured in a 37C shaking incubator. After 18h of incubation, 1mL of the OP50 culture was pelleted, then resuspended in 40uL of M9. This was used as the mounting buffer. Before each recording, we made a thin, flat agar pad (2.5cm x 1.8cm x 0.8mm) with NGM containing 2% agar. On the 4 corners of the agar pad, we placed a single layer of microbeads with a diameter of 80um to alleviate the pressure of the coverslip on the worm. Then a worm was picked to the middle of the agar pad, and 9.5uL of the mounting buffer was added on top of the animal. Finally, a glass coverslip (#1.5) was added on top of the worm. This caused the mounting buffer to spread evenly across the slide. We waited for 5 minutes after mounting the animal before imaging.

#### Procedure for NeuroPAL imaging

For NeuroPAL recordings, animals were imaged as described above, but they were subsequently immobilized by cooling, after which multi-spectral information was captured. The slide was mounted back on the confocal with a thermo-electric cooling element attached to it, set to cool the agar temperature to 4°C (Wang et al., 2022). A closed-loop temperature controller (TEC200C, Thorlabs) with a micro-thermistor (SC30F103A, Amphenol) embedded in the agar kept the agar temperature at the 1 °C set point. Once the temperature reached the set point, we waited 5 minutes for the worm to be fully immobilized before imaging. Details on exactly which multi-spectral images were collected are in the NeuroPAL annotation section below.

#### Heat stimulation

For experiments involving heat stimulation, animals were recorded using the procedure described above, but were stimulated with a 1436-nm 500-mW laser (BL1436-PAG500, Thorlabs) a single time 4 or 6 min into the recording. The laser was controlled by a driver (LDC220C, Thorlabs) and cooled by the built-in TEC controller and a temperature controller (TED200C, Thorlabs). The light emitted by the laser fiber was collimated by a collimator (CFC8-C, Thorlabs) and expanded to be about 600 um at the sample plane. The laser light was fed into the NIR brightfield path via a dichroic with 1180-nm cutoff (DMSP1180R, Thorlabs). We determined the amplitude and kinetics of the heat stimulus in calibration experiments where temperature was determined based on the relative intensities of rhodamine 110 (temperature-insensitive) and rhodamine B (temperature-sensitive). This procedure was necessary because the thermistor size was considerably larger than the 1436-nm illumination spot, so it could not provide a precise measurement of temperature within the spot. Slides exactly matching our worm imaging slides were prepared with dyes added (and without worms). Dyes were suspended in water at 500mg/L and diluted into both agar and mounting buffer at a 1:100 dilution (final concentration of 5mg/L). Rhodamine 110 was imaged using a 510/20 bandpass filter and rhodamine B was imaged was 610LP filter. We recorded data using the confocal light path during a calibration procedure where a heating element ramped the temperature of the entire agar pad from room temp to >50°C. Temperature was simultaneously recorded via a thermistor embedded on the surface of the agar, approximating the position of the worm. Fluorescence was also recorded at the same time, at the precise position where the worm’s head is imaged. This yielded a calibration curve that mapped the ratio of Rhodamine B/Rhodamine 110 intensity at the site of the worm’s head onto precise temperatures. Slides were then stimulated with the 1436-nm laser using identical setting to the experiments with animals. The response profile of the ratio of the fluorescent dyes was then converted to temperature. We quantitatively characterized the change in temperature, noting the mean temperature over the first second of stimulation (set to be exactly 10.0°C) and its decay (0.39 sec exponential decay rate, such that it returns to baseline within 3sec).

### Extraction of behavioral parameters from NIR videos

We quantified behavioral parameters of recorded animals by analyzing the NIR brightfield recordings. All of these behaviors are initially computed at the NIR frame rate of 20Hz, and then transformed into the confocal time frame using camera timestamps, averaging together all of the NIR data corresponding to each confocal frame.

#### Velocity

First, we read out the (x,y) position of the stage (in mm) as it tracks the worm. To account for any delay between the worm’s motion and stage tracking, at each time point we added the distance from the center of the image (corresponding to the stage position) to the position of the metacorpus of pharynx (detected from our neural network used in tracking). This then gave us the position of the metacorpus over time. To decrease the noise level (eg: from neural network and stage jitter), we then applied a Group Sparse Total Variation Denoising algorithm to the metacorpus position. Differentiating the metacorpus position then gives us a movement vector of the animal.

Because this movement vector was computed from the location of the metacorpus, it contains two components of movement: the animal’s velocity in its direction of motion, and oscillations of the animal’s head perpendicular to that direction. To filter out these oscillations, we projected the movement vector onto the animal’s facing direction, i.e. the vector from the grinder of the pharynx to its metacorpus (computed from the stage-tracking neural network output). The result of this projection is a signed scalar, which is reported as the animal’s velocity.

#### Worm spline and body angle computation

To generate curvature variables, we trained a 2D U-Net to detect the worm from the NIR images. Specifically, this network performs semantic segmentation to mark the pixels that correspond to the worm. To ensure consistent results if the worm intersects itself (for instance, during an omega-turn), we use information from worm postures at recent timepoints to compute where a self-intersection occurred, and mask it out. Next, we compute the medial axis of the segmented and masked image and fit a spline to it. Since the tracking neural network was more accurate at detecting the exact position of the worm’s nose, we set the first point of the spline to the point closest to the tracking neural network’s nose position. We next compute a set of points along the worm’s spline with consistent spacing (8.85 μm along the spline) across time points, with the first point at the first position on the spline. Body angles are computed as the angles that vectors 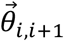 between adjacent points make with the *x*-axis; for example, the first body angle would be the angle that the vector 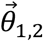 between the first and second point makes with the *x*-axis, the second body angle would be 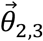, and so on.

#### Head curvature

Head curvature is computed as the angle between the points 1, 5, and 8 (ie: the angle between 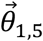 and 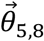). These points are 0 μm, 35.4 μm, and 61.9 μm along the worm’s spline, respectively.

#### Angular velocity

Angular velocity is computed as smoothed 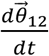, which is computed with a linear Savitzky-Golay filter with a width of 300 time points (15 seconds) centered on the current time point.

#### Body curvature

Body curvature is computed as the standard deviation of 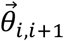 for *i* between 1 and 31 (ie: going up to 265 μm along the worm’s spline). This value was selected such that this length of the animal would almost never be cropped out of the NIR camera’s field of view. To ensure that these angles are continuous in *i*, they may each have 2*π* added or subtracted as appropriate.

#### Feeding (pumping rate)

Pumping rate was manually annotated using Datavyu, by counting each pumping stroke while watching videos slowed down the 25% of their real-time speeds. The rate is then filtered via a moving average with a width of 80 time points (4 seconds) to smoothen the trace into a pumping rate rather than individual pumping strokes.

### Extraction of normalized GCaMP traces from confocal images

We developed the Automatic Neuron Tracking System for Unconstrained Nematodes (ANTSUN) software pipeline to extract neural activity (normalized GCaMP) from the confocal data consisting of a time series of z-stacks of two channels (TagRFP-T or mNeptune2.5 for the marker channel and GCaMP7f for the neural activity channel). Each processing step is outlined below.

#### Pre-processing

The raw images first go through several pre-processing steps before registration and trace extraction. For datasets with a gap in the middle, all of the following processing is done separately and independently on each half of the dataset.

##### Shear correction

Shear correction is performed on the marker channel, and the same parameters are also used to transform the activity channel. Since the images in a z-stack are acquired over time, there exists some translation across the images within the same z-stack, causing some shearing. To resolve this, we wrote a custom GPU accelerated version of the FFT based subpixel alignment algorithm (Guizar-Sicairos et al., 2008). Using the alignment algorithm, each successive image pair is aligned with x/y-axis translations.

##### Image cropping

We crop the z-stacks to remove the irrelevant non-neuron pixels. For each z-stack in the time series, the shear-corrected stack is first binarized by thresholding intensity. Using principal component analysis on the binarized worm pixels, the rotation angle about the z-axis is determined. Then the stack is rotated about the z-axis with the determined angle to align the worm’s head. Then the 3D bounding box is determined using the list of worm pixels after the rotation. Finally, the rotated z-stack is cropped using the determined 3D bounding box. Similar to shear correction, this procedure is first done on the marker channel, and the same parameters are then applied to the activity channel.

##### Image filtering using total variation minimization

To filter out noise on the marker channel images, we wrote a custom GPU accelerated version of the total variation minimization filtering method, commonly known as the ROF model (Rudin et al., 1992). This method excels at filtering out noises while preserving the sharp edges in the images. Note that the activity channel is kept unfiltered for GCaMP extraction.

#### Registering volumes across time points

To match the neurons across the time series, we register the processed z-stacks across time points. However, simply registering all time points to a single fixed time point is intractable because of the high amount of both global and small-scale deformations. To resolve this, we compute a similarity metric across all possible time point pairs that reports the similarity of worm postures. We then use this metric to construct a registration graph where nodes are timepoints and edges are added between timepoints with high posture similarity. The graph is constrained to be fully connected with an average connectedness of 10. Therefore, it is possible to fully link each time point to every other time point. Using this graph, we register strategically chosen pairs of z-stacks from different time points (i.e. the ones with edges). The details of the procedure are outlined below.

##### Posture similarity determination

For each z-stack, we first find the anterior tip of the animal using a custom trained 2D U-Net, which outputs the probability map of the anterior tip given a maximum intensity projection of the z-stack. We then fit a spline across the centerline of the neuron pixels beginning at the determined anterior tip, which is the centroid of the U-Net prediction. Using the spline, we compare across time points pairs to determine the similarity.

##### Image registration graph construction

Next, we construct a graph of registration problems, with time points as vertices. For each time point, an edge is added to the graph between that time point and each of the ten time points with highest similarity to it. The graph is then checked for being connected.

##### Image registration

For each registration problem from the graph, we perform a series of registrations that align the volumes, iteratively in multiple steps in increasing complexity: Euler (rotation and translation), affine (linear deformation), and B-spline (non-linear deformation). In particular, the B-spline registration is performed in three scales, decreasing from global (the control points are farther apart) to local (the control points are placed closer together) registration. The image registrations and transformations are performed using elastix on OpenMind, a high-performance computing cluster. They are performed on the mNeptune2.5 marker channel.

#### Channel alignment registration

To align the two cameras used to acquire the marker and the activity channels, we perform Euler (translation and rotation) registration across the two channels over all time points. Then we average the determined transformation parameters from the different time points and apply across all time points.

#### Neuron ROI determination

To segment out the pixels and find the neuron ROIs, we first use a custom trained 3D U-Net. The instance segmentation results from the U-Net are further refined with the watershed algorithm.

##### Simultaneous semantic and instance segmentation with 3D U-Net

We trained a 3D U-Net to simultaneously perform semantic and instance segmentation of the neuronal ROIs in the z-stacks of the unfiltered marker images. To achieve instance segmentation, we labeled and assigned high weights to the boundary pixels of the neurons, which guides the network to learn to segment out the boundaries and separate out neighboring neurons. Given a z-stack, the network outputs the probability of each pixel being a neuron. We threshold and binarize this probability volume to mark pixels that are neurons.

##### Instance segmentation refinement

To refine the instance segmentation results from the 3D U-Net, we perform instance segmentation using the watershed algorithm. This generates, for each time point, a set of ROIs in the marker image corresponding to distinct neurons.

#### Neural trace extraction

##### ROI Similarity Matrix

To link neurons over time, we first create a symmetric *N × N* similarity matrix, where *N* is the number of total ROIs detected by our instance segmentation algorithm across all time points. Thus, for each index *i* ∈ 1:*N* in this matrix, we can define the corresponding time point *t_i_* and the corresponding ROI *r_i_* from that time point. This matrix is sparse, as its (*i,j*)*th* entry is nonzero only if there was a registration between *t_i_* and *t_j_* that maps the ROI *r_i_* to *r_j_*. In the case of such a registration existing, the (*i,j*)*th* entry of the matrix is set to a heuristic intended to estimate confidence that the ROIs *r_i_* and *r_j_* are actually the same neuron at different timepoints. This heuristic includes information about the quality of the registration mapping *r_i_* to *r_j_* (computed using Normalized Correlation Coefficient), the fractional volume of overlap between the registration-mapped *r_i_* and *r_j_* (i.e. position similarity), the difference in marker expression between *r_i_* and *r_j_* (i.e. similarity of mNeptune expression), and the fractional difference in volume between *r_i_* and *r_j_* (i.e. size similarity of ROIs). The diagonal of the matrix is additionally set to a nonzero value.

##### Clustering the Similarity Matrix

Next, we cluster the rows of this similarity matrix using a custom clustering method; each resulting cluster then corresponds to a neuron. First, we construct a distance matrix between rows of the similarity matrix using L2 Euclidean distance. Next, we apply minimum linkage hierarchical clustering to this distance matrix, except that after a merge is proposed, the resulting cluster is checked for ROIs belonging to the same time point. If too many ROIs in the resulting cluster belong to the same time point, that would signify an incorrect merge, since neurons should not have multiple different ROIs at the same time point. Thus, if that happens, the algorithm does not apply that merge, and continues with the next-best merge. This continues until the algorithm’s next best merge reaches a merge quality threshold, at which point it is terminated, and the clusters are returned. These clusters define the grouping of ROIs into neurons.

##### Linking multiple datasets

For datasets that were recorded with a gap in the middle, the above process was performed separately on each half of the data. Then, the above process was repeated to link the two halves of the data together, except that only two edges that must connect to the other half of the data are added to the registration graph per time point, and the clustering algorithm does not merge clusters beyond size 2.

##### Trace extraction

Next, neural traces are extracted from each ROI in each time point belonging to that neuron’s cluster. Specifically, we obtain the mean of the pixels in the ROI at that time point. This is done in both the marker and activity channels. They are then put through the following series of processing steps:

- Background-subtraction, using the median background per channel per time point.
- Deletion of neurons with too low of signal in the activity channel (mean activity lower than the background), or too few ROIs corresponding to them (less than half of the total number of time points).
- Correction to account for laser intensity changing halfway through our recording sessions (done separately on each channel based on intensity calibration measurements taken at various values of laser power).
- Linear interpolation to any time point that lacked an ROI in the cluster.
- Division of the activity channel traces by the marker channel traces, to filter out various types of motion artifacts. These divided traces are the neural activity traces.

##### Bleach correction

We then compute the mean neural activity (averaged across all neurons) over the entire time range, and fit a one-parameter exponential bleaching model to it. The bleaching model was initialized such that it had value equal to the median neural activity value at the median time point, and it was fit using log-MSE error to the averaged neural activity value. A small number of datasets with very high bleaching (determined using the exponential decay parameter of the bleaching model) were excluded from all analysis. Each neural activity trace is then divided by the best-fit bleaching curve; the resulting traces are referred to as *F*. In our SWF360 analysis, we refer directly to *F;* the trace heatmaps shown in this paper are 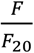 (where *F*_20_ is the 20^th^ percentile, computed separately for each neuron); we also display z-scored neural activity in many figure panels, as indicated; and the CePNEM models are fit by z-scoring each neuron separately.

##### Controls to test whether neurons are correctly linked over time

We ran a control to test whether neurons were being mismatched by our registration process. We did this by processing data from our SWF360 strain that expresses GFP at different levels in different neurons (*eat-4::NLS-GFP).* The recording was made with a gap and was processed identically to GCaMP datasets with gaps in the middle, thus also serving as a test of inter-gap registration. This SWF360 recording allows us to detect errors in neuron registration, since GFP-negative neuron could briefly become GFP-positive or vice versa. We quantified this by setting a threshold of median(*F*) > 1.5 to call a neuron a GFP neuron. This threshold resulted in Frac_GFP_ = 27% of neurons being quantified as containing GFP, which is about what was expected for the promotors expressed. Then, for each neuron, we quantified the number of time points such that the neuron’s activity F at that time point differed from its median by more than 1.5, and exactly one of [the neuron’s activity at that time point] and [its median activity] was larger than 1.5. These time points represent mismatches, since they correspond to GFP-negative neurons that were mismatched to GFP-positive neurons (if the neuron’s activity increased at the time point) or vice versa (if its activity decreased). We then computed an error rate of 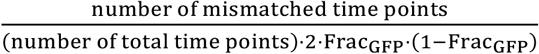 as an estimate of the mis-registration rate of our pipeline. The 2 · Frac_GFP_ · (1 – Frac_GFP_) term corrects for the fact that mis-registration errors that send GFP-negative to other GFP-negative neurons, or GFP-positive to other GFP-positive neurons, would otherwise not be detected by this analysis. This error rate came out to 0.3%, so we conclude that artifacts resulting from mismatched neurons are a negligible component of our data.

### Annotation of neural identities using NeuroPAL

The identities of neurons were determined via NeuroPAL using the following procedure. We obtained a series of images from each recorded animal, while the animal was immobilized after the freely-moving GCaMP recording (recording and immobilization procedures described above):

(1-3) Spectrally isolated images of mTagBFP2, CyOFP1, and mNeptune2.5. We excited CyOFP1 using the 488nm laser at 32% intensity under a 585/40 bandpass filter. mNeptune2.5 was recorded next using a 637nm laser at 48% intensity under a 655LP-TRF filter, in order to not contaminate this recording with TagRFP-T emission. Finally, mTagBFP2 was isolated using a 405nm laser at 27% intensity under a 447/60 bandpass filter.

(4) An image with TagRFP-T, CyOFP1, and mNeptune2.5 (all of the “red” markers) in one channel, and GCaMP7f in the other channel. As described below, this image was used for neuronal segmentation and registration with both the freely moving recording and individually isolated marker images. We excited TagRFP-T and mNeptune2.5 via 561nm laser at 15% intensity and CyOFP1 and GCaMP6f via 488nm laser at 17% intensity. TagRFP-T, mNeptune2.5, and CyOFP1 were imaged with a 570LP filter and GCaMP6f was isolated using a 525/50 bandpass filter.

All isolated images were recorded for 60 timepoints. We increased the signal to noise ratio for each of the images by first registering all timepoints within a recording to one another and then averaging the transformed images. Finally, we created the composite, 3-dimensional RGB image by setting the mTagBFP2 image to blue, CyOFP1 image to green, and mNeptune2.5 image to red as done by Yemini et al. (2021) and manually adjusting the intensity of each channel to optimally match their manual.

The neuron segmentation U-Net was run on the “all red” image and we then determined the identities of U-net identified neurons using the NeuroPAL instructions. The landmarks in the NeuroPAL atlas were identified first and the identities of the remaining neurons were subsequently determined by comparing the individual channel intensities, overall coloring, and relative positioning of the cells. In some cases, neuronal identities could not be determined with certainty due a number of factors including: unexpectedly dim expression of one or more fluorophores, unexpected expression of a fluorophore in cells not stated to express a given marker, and extra cells in a region expressing similar intensities when no other cells are expected. Rarely, multiple cells were labeled as potential candidates for a given neuron and the most likely candidate (based on position, coloring, and marker intensity) was used for analysis. If a cell was not bright enough to be distinguished from its neighbors or was undetected by the neuron segmentation U-Net, we left it unlabeled.

Finally, the neural identity labels from the RGB image were mapped back to the GCaMP traces from the freely-moving animal by first registering each fluorophore-isolated image to the image containing all of the red markers. The “all red” image was then registered back to the freely moving recording, permitting mapping of neuronal labels back to GCaMP traces.

### Decoding behavior from neural activity

#### Full activity, current behavior

We trained L1-regularized linear decoder models to predict the worm’s current velocity, head curvature, feeding rate, angular velocity, and body curvature based on its current (z-scored) neural activity. To set the regularization parameter, we withheld three datasets that were randomly selected from the set of datasets with feeding standard deviation of at least 0.5. The other eleven datasets were used to evaluate decoder performance. The decoders were evaluated using five-fold cross-validation splits. All behaviors were z-scored for the decoder, and the decoder accuracy is reported as one minus the MSE between the decoder’s prediction and actual behavior, evaluated on the test-time data.

#### Model residuals, current behavior

We computed model residuals for each neuron by taking that neuron’s activity and subtracting the modeled n[t] (computed based off of the median of all posterior CePNEM parameters for that neuron), and then z-scoring the resulting residual trace. We then trained separate decoder models using the same procedure as above, except using the model residuals instead of neural activity. We regularized these decoders separately using the same three set-aside datasets.

#### Decoding past behavior (Figure 2D)

We trained linear decoder models to predict the average velocity of the worm at time points in the past, based on the worm’s current (z-scored) neural activity; only neurons that encoded velocity were included. The models were trained on data from all 14 SWF415 animals. A separate model was trained for each time point in the past. The average velocity was computed in the window spanning (Δt — 8, Δt + 8] where Δt is the number of time points into the past (Δt = 0 is current). This approximately corresponds to a 10-sec time window. Velocity across the full 1600 time points was z-scored before being averaged. Each dataset was split into 5 segments for cross-validation, with 100-timepoint gaps in between to prevent the training time information from spilling over to the test time segment. The models were regularized using an elastic net (L1 and L2).

As a control, separate models were trained that attempted to predict shifted velocity, which should scramble the relationship between neural activity and behavior. Velocity was circularly shifted by an amount between 125 and 600 time points, and, additionally, shifts that would result in a correlation of greater than 0.2 with actual velocity were discarded. 50 such decoders were trained, each using a different, randomly-selected shift. The performance of the decoder trained to predict averaged velocity Δt time points into the past was then defined as the difference between the cost (square root of MSE) of that decoder and the average cost of each of the 50 decoders trained on shifted velocity.

To ensure that decoder performance based on neural activity with Δt > 0 was actually a representation of historical velocity information, and not simply due to the autocorrelative nature of velocity, a separate family of decoders were trained that was given the worm’s current (z-scored) velocity as input instead of neural activity. The error of those decoders to their shifted controls is also displayed in Figure 2D. Finally, to estimate the maximum possible performance of these decoder models, separate “perfect” decoders were trained that were given the worm’s (z-scored) velocity at time points t + Δt for each value of Δt ∈ (—8,108), and were then subjected to the same shift test.

#### *C. elegans* Probabilistic Neural Encoding Model (CePNEM)

##### Fitting procedure

###### Overview of fitting approach

Let N be a neural trace from an animal, B be the behaviors of that animal, and X be the model parameters that we are trying to fit. Then the goal our model fitting procedure is to estimate the probability distribution of model parameters given our observations, namely P(X|N, B). Our model defines the likelihood P(N|X, B) – that is, the likelihood of observing a set of neural data given a set of model parameters and behavioral data. Our prior distributions define P(X|B); in this case, our prior distributions on model parameters are independent of the animal’s behaviors, so P(X|B) = P(X). Therefore, by Bayes’ rule,

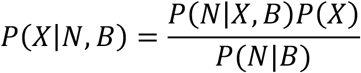

Unfortunately, *P*(*N*|*B*) is difficult to compute. Crucially, however, it does not depend on the model parameters *X*. This means that by comparing the value of *P*(*N*|*X, B*)*P*(*X*) for different values of *X*, we can make meaningful insights into the distribution of *P*(*X*|*N, B*). In particular, we can define a Markov chain that defines a sequence of *X_t_*, where *X*_*t*+1_ is a stochastic “proposal function” of *X_t_*. The idea is that the proposal function can be biased to walk toward regions in parameter space with higher likelihood. Indeed, there are a family of algorithms, such as Metropolis-Hastings (Hastings, 1970) and Hamiltonian Monte Carlo (Neal, 2011) that define such proposal functions. In particular, the proposal functions defined by these algorithms have the property that, in the limit of generating an infinitely long Markov chain, sampling from the chain is equivalent to sampling from the true posterior distribution *P*(*X*|*N, B*). A description of the noise model and priors used are below.

###### MCMC fitting procedure

Of course, in practice, we do not have computational resources for an infinitely long chain, so it is necessary to ensure that the chain can replicate the posterior distribution in a manageable amount of time. To accomplish this, we use a mixture of Metropolis-Hastings (MH) and Hamiltonian Monte Carlo (HMC) steps. The HMC step uses gradient information and tries to move the chain towards regions of higher likelihood. The other MH steps are intended to help the chain get out of local optima by using information about the structure of the model, so the Markov chain can better explore the full parameter space. Specifically, one iteration of our fitting algorithm involves the following steps (here 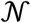 is once again the normal distribution, and 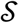 is drawn uniformly at random from the set [-1,1]), and i is the current iteration of the algorithm:

- MH proposal: 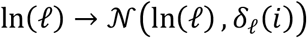
- MH proposal: 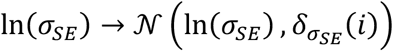
- MH proposal: 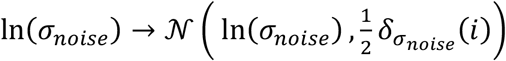
- HMC proposal on parameters *c_vT_, c_v_, c_θh_, c_p_,b,n* (0) ln(*s*) with *ϵ* = *δ_HMC_* (*i*)
- MH proposal: 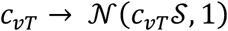
- MH proposal (note that the instances of 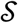 are drawn independently):

- 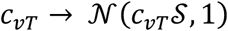
- 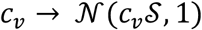
- 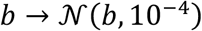

After each iteration of the algorithm, the learning rate parameters *δ* are updated as follows: If the respective proposal was accepted, its *δ* parameter is multiplied by 1.1; otherwise, it is divided by 1.1. (They are all initialized to 1.) In this fashion, the learning rate will converge to a value such that about half of the proposals will be accepted, resulting in a faster overall convergence of the MCMC chain. To construct the posterior samples used in our analysis, we run this MCMC chain for 11,000 iterations, and discard the first 1,000 (including the initialization point). The remaining 10,001 points in the posterior distribution are referred to as particles.

###### MCMC chain initialization

Despite our efforts to use MH proposal steps to prevent the MCMC procedure from falling into local optima, we found that the algorithm still occasionally got stuck, preventing it from finding a good approximation to the true posterior. To remedy this, we added a Likelihood Weighting initialization step consisting of sampling 100,000 points from the prior distribution of model parameters and selecting the point with the highest likelihood under our model, given the neural and behavioral data to be fit. This point is then used to initialize the MCMC chain detailed above. Using Gen allowed us to combine Likelihood Weighting with a custom set of MCMC kernels, described above. This can be viewed as a form of resample-move SMC (Berzuini and Gilks, 2001). Gen also allowed us to automate validation via simulation-based calibration (see next section). These capabilities are not provided by other probabilistic programming languages such as Stan.

##### Simulation-based calibration

To ensure that our fitting process gave a calibrated description of the true model posterior, we performed simulation-based calibration (Talts et al., 2020). In this procedure, we generated 4,000 sample traces from the model distribution *P*(*X, N* |*B*) using the prior distribution for X. 500 traces were generated using each of eight total values of B: two 800-time-point subsegments from each of four animals (two SWF415, and two SWF702 animals). We then ran our MCMC inference procedure on each sample (three of the 4,000 traces timed out and were discarded). After fitting, we then compared the sampled posterior distribution from our inference algorithm to the ground-truth parameter values using a rank test with 128 bins. If our inference process was giving unbiased estimates of the posterior distribution, then across all of our traces, the distribution of these ranks should be the uniform distribution.

We used a *χ*^2^ test to differentiate the observed ranks from the uniform distribution, and found that 9 of the 10 model parameters passed the test at p=0.05. The final parameter, the EWMA decay constant s, seemed to have a minor bias towards larger values, meaning that our fitting algorithm is prone to occasionally overestimate this parameter. However, we quantified an upper bound on the degree of this overestimation by computing the maximum deviation of the CDF of the observed rank distribution for s, compared with the predicted CDF from the uniform distribution, and found a value of 3.5%. This means that the fits of at most 3.5% of encoding neurons will be affected by this minor bias, which is less than an average of 4 per animal. Thus, we do not believe this minor bias will substantially affect the results described in this paper.

#### CePNEM Noise Model

The CePNEM model uses a Gaussian process noise model adding together a white-noise kernel and a squared exponential kernel. The white-noise kernel is intended to capture measurement noise in our neural data, which is expected to be independent between time points, while the squared exponential kernel is intended to capture variance in neural activity unrelated to behavior, which may have a slower timescale. The squared-exponential noise term is critically important, as otherwise the model will be forced to try to explain all autocorrelation in neural activity with behavioral information, resulting in severe overfitting.

The white-noise kernel *K_GN_* has standard deviation *σ_noise_* and thus its covariance matrix is 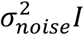. The squared-exponential kernel *K_SE_* has standard deviation *σ_SE_* and length scale *ℓ*, giving a covariance matrix defined by 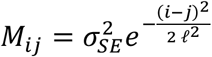. The full noise model is then the Gaussian process model with kernel *K_GN_* + *K_SE_*, which is then added to the timeseries of the rest of the model fit to generate the likelihood of a given neural activity trace under CePNEM.

#### CePNEM Prior Distributions

Here 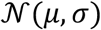 is the normal distribution with mean *μ* and standard deviation *σ*. Since the neural traces being fit are all z-scored, the priors on the behavioral parameters are also standardized. The prior on the moving average term *s* was based on preliminary data from fitting previous, conventional versions of our model. The priors on the noise terms were intended to be wide enough to accommodate both neurons that are well-explained by behaviors (in which case, the model would assign them a low noise value), and neurons that contain little to no information about behaviors (in which case, the model would assign them a high noise value).

### Statistical tests to determine encoding properties of neurons

#### Deconvolved activity matrix

In order to be able to make statistical assertions about the neural encoding of behavior based on the posterior distributions from CePNEM fits, we first needed to transform model parameters into a more intuitive space. To accomplish this, for each neuron, we constructed a 10001 × 4 × 2 × 2 deconvolved activity matrix *M* constructed as follows: *M_nijk_* corresponds to the modeled activity of the nth particle from that neuron’s CePNEM fit at velocity *V*[*i*], head curvature *θH*[*j*], and pumping rate *P*[*k*]. Here, where *θh* is the animal’s head curvature (dorsal is positive) and *μ* is the animal’s pumping rate over the course of the recording, we have:

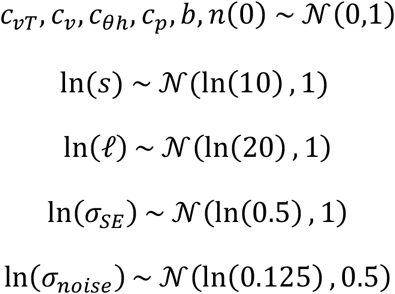

For this calculation, the EWMA and noise components are excluded from the modeled activity; the idea is that this matrix contains information about the neuron’s activity at high and low values of each behavior, so we can now run analyses on this matrix and not have to take into account the actual behavior of the animal. In particular, many simple combinations of entries in this matrix have intuitive meanings:

- The slope of the neuron’s tuning to velocity during forward locomotion is

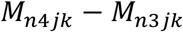
- The slope of the neuron’s tuning to velocity during reverse locomotion is

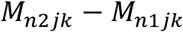
- The neuron’s deconvolved forwardness (overall slope of the neuron’s tuning to velocity) is

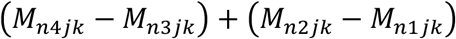
- The rectification of the neuron’s tuning to velocity (difference between forward and reverse slopes) is

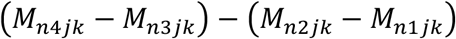
- The slope of the neuron’s tuning to head curvature during forward locomotion (positive means dorsal during forward) is

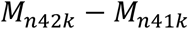
- The slope of the neuron’s tuning to head curvature during reverse locomotion (positive means dorsal during reverse) is

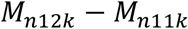
- The neuron’s deconvolved dorsalness (overall slope of the neuron’s tuning to head curvature) is

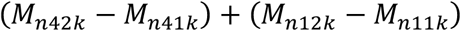
- The rectification of the neuron’s tuning to head curvature with respect to locomotion direction (positive means the neuron is more dorsal during forward; negative means the neuron is more ventral during forward) is

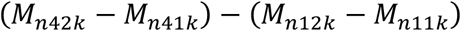
- The neuron’s tuning to feeding follows the same pattern as its tuning to head curvature.

Importantly, the linear structure of the multiplexing component of CePNEM implies that the value of the unset parameters *i,j, k* in the expressions above do not change the value of those expressions. For head curvature, since worms can lay on either side, we manually checked the location of the animal’s vulva from the NIR recordings of each animal and flipped dorsal/ventral labels as appropriate.

#### Statistical encoding tests

With the intuition derived from the deconvolved activity matrix, for each particle in the posterior distribution of the neuron, we can ask whether that particle satisfies a certain property. For example, to categorize a particle as representing forward locomotion, we would check whether that particle had a sufficiently large deconvolved forwardness value. Specifically, we would check whether its deconvolved forwardness value was at least max (*ξ*_1_, *ξ*_2_), where 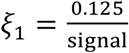 (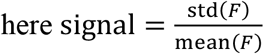 and *F* is the un-normalized ratiometric fluorescence of the neuron in question), and 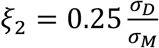 (here *σ_D_* is the standard deviation of the model fit corresponding to that particle with *s* = 0 and *σ_M_* is the standard deviation of the model fit corresponding to that particle). The number 0.125 was selected based on its ability to filter out the small amount of motion artifacts observed in our three GFP control datasets (see Methods section on that control below). Specifically, we chose a value that filtered out almost all of the motion artifacts (leaving only 2.1% of GFP neurons showing false behavioral encoding), while removing as few true encodings from our GCaMP data as possible. Similarly, the number 0.25 was selected based on its ability to filter out extremely weak correlations between neural activity and behavior, which was measured by our controls attempting to fit neurons with behaviors from different animals (after the correction, only 2.7% of such neurons showed behavioral encoding). The 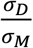 term is a correction for the fact that neurons with large *s* values will have higher values in *M*. If the particle’s deconvolved forwardness value was at least max (*ξ*_1_, *ξ*_2_), it would be classified as representing forward locomotion.

By the same token, we would classify a particle as representing reverse locomotion if its deconvolved reverseness (negative forwardness) value was at least max (*ξ*_1_, *ξ*_2_), we would classify a particle as representing more dorsal information during forward locomotion if its rectification to head curvature with respect to locomotion direction was at least max (*ξ*_1_, *ξ*_2_), and so on.

Now that we can classify particles, we can create empirical p-values for neurons based on the fraction of their particles that share a category. For example, if 98% of particles for a neuron are classified as representing forward locomotion, then that neuron’s *p*-value for forward locomotion would be 0.02. We can then construct a list of such *p* values computed for each neuron in an animal that was fit with CePNEM and use Benjamini-Hochberg correction with FDR=0.05 to get a list of forward-encoding neurons in that animal. We can similarly get a list of reversal neurons, dorsally-rectified head curvature neurons, neurons activated by feeding during forward locomotion (i.e. have a positive slope to feeding during forward locomotion), and so on.

To construct larger categories, such as neurons with any behavioral encoding, or neurons with head curvature encoding, another multiple hypothesis correction step is necessary. For this step, we first use Bonferroni correction on opposing categories where it is impossible for a neuron to have both categories (for instance, dorsal and ventral tuning), followed by a Benjamini-Hochberg correction step on the Bonferroni-corrected *p*-values. We then proceed with the inter-neuron Benjamini-Hochberg correction, as before.

A neuron is categorized as encoding head curvature if it expresses statistically significant information about any of the four head curvature categories outlined above, in either direction; feeding encoding is computed similarly. A neuron is categorized as encoding velocity if it either expresses statistically significant information about any of the four velocity categories, or if it expresses statistically significant information about any of the rectified categories, since rectification of head curvature or feeding based on forward/reverse locomotion state is a form of velocity information. A neuron is categorized as encoding if it has statistically significant information in any of the tests. Note that for datasets without any feeding information (defined as the 25^th^ and 75^th^ percentile of feeding in that dataset being the same, causing *P*[1] = *P*[2]), neurons cannot encode feeding information, so feeding is not included in the multiple-hypothesis correction to check whether a neuron encoded any behavior.

#### Forwardness, Dorsalness, and Feedingness

The forwardness metric for a neuron is computed as the median of 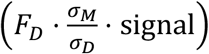 over all particles for that neuron, where *F_D_* is the deconvolved forwardness of that particle, and *σ_M_*, *σ_D_*, and signal are as before. Dorsalness and feedingness are computed in a similar fashion.

#### Time ranges

One final note is that all neurons are fit twice – once over the first half of the data, and once over the second half. This is done because a large number of our SWF415 datasets have a gap in the middle, and due to the EWMA term in our model, it would be difficult to fit a model in a time range that included a gap. Thus, for consistency between all our datasets, we fit all of our SWF415 and NeuroPAL datasets in this manner.

For Figure 2A, the encoding statistics are computed on a per-neuron basis, with an additional Benjamini-Hochberg correction step to account for the fact that each neuron got two chances to qualify as encoding. Time ranges with insufficient feeding variance (this time, defined as the difference between the 25^th^ and 75^th^ percentile of feeding being at most 0.5) are excluded from feeding analysis. To avoid different behaviors having different amounts of available data, animals that never had sufficient feeding variance are excluded from Figure 2A entirely. For Figure 2B, the same analysis is used, and there is an additional multiple-hypothesis step across the three behaviors. For Figures 2C and S2G, all time ranges are used. Fits on different time ranges from the same animal are added to the CDF independently of each other, but only encoding neurons are included. For example, a neuron that encoded behavior in both time ranges would have its EWMA timescale from both fits added to the CDF, while a neuron that only encoded behavior once would have that EWMA timescale added. In Figure S2G, only neurons that statistically significantly encoded forward or reverse locomotion are included.

#### Neuron Subcategorization

We next sought to combine various pieces of information from our encoding analysis together to generate a holistic view of how a given neuron is tuned to a given behavioral parameter. To accomplish this, we sorted neurons as follows (this analysis is done independently on each time range):

- If the neuron had a different sign to its tuning to behavior during forward and reverse (eg: a slow neuron that has a positive slope in its tuning to velocity during reversal, but a negative slope during forward locomotion), then the neuron was categorized as such. In Figures 2G–2I, these neurons would appear in the bins (+,-) and (-,+); for head curvature, they would be (D,V) or (V,D).
- Otherwise, if the neuron has rectified tuning to the behavior (depending on the behavior, one of the following categories: forward slope > reverse slope, reverse slope < forward slope, more dorsal during forward, more ventral during more activated during forward, more activated during forward, or more inhibited during forward), it will be placed in one of the four rectified bins (+,0), (-,0), (0,-), or (0,+), depending on the sign of the rectification and sign of the slopes of the neural tuning to behavior.
- Otherwise, if the neuron had the same slope during both forward and reverse movement, it will be classified in one of the two analog bins (+,+) or (-,-) depending on the sign of that slope. Notably, it would be placed in a rectified bin (and not an analog bin) if it had rectified information, even if it had the same slope during both forward and reverse locomotion.
- If none of the above were true, the neuron lacked statistical significance in at least two of the three parameters (forward slope, reversal slope, rectification) with respect to the behavior in question, and it will be excluded from Figures 2E–2G.

In Figures 2H and 2I, the neurons had the same tuning in both time ranges, and the EWMA values reported incorporate fits from both time ranges.

### Median model fits

For display purposes, or analyses where it was necessary to represent a neuron with a single model, we computed the median model by computing *n_i_* [*t*] for each set of parameters *i* in the neuron’s posterior distribution, and then defining *n_med_* [*t*] = median_*i*_ (*n_i_* [*t*]). This is what is meant by “median CePNEM fit” unless otherwise noted.

### Encoding strength

Encoding strength is a metric designed to approximate the information content a neuron contains about each behavior, given its CePNEM model fits. It is computed by generating three model traces *n_v_*, *n_θh_*, and *n_P_*, each of which is identical to the full model *n*[*t*] except that the behavior *i* is set to 0 for model *n_i_*. Thus, the MSE between *n* and *n_i_* provides a metric of how important behavior i was for the neuron. We compute the encoding strength of a neuron to behavior *i* as the ratio 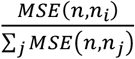.

### GFP Control

We wanted to ensure that we would not spuriously detect motion artifacts as encodings of behavior. To do this, we used our pan-neuronal GFP control line SWF467, which by definition should not have any neurons detect as encoding behavior. We fit our GFP datasets with CePNEM and applied the same encoding analysis to this strain and found that only 2.1% of neurons showed behavioral encoding, compared with 58.6% in the SWF415 strain, suggesting that the vast majority (>95%) of our detected encodings are not due to motion artifacts.

### Scrambled Control

We furthermore wanted to ensure that the model would not overfit to spurious correlations between neural activity and behavior. To accomplish this, we fit 11 SWF415 animals with CePNEM, but replaced the behaviors *v, θh*, and *p* with spurious behaviors from other recorded animals, which should result in few neurons showing behavioral encoding. The spurious behaviors were generated as follows: we first assign pairs of datasets to minimize the behavioral correlation across the datasets within a given pair. To do this, we compute correlation across all possible behavior and dataset combinations. After that, we determine the pairing such that it minimizes the maximum absolute cross-correlation value across all pairings. To penalize high correlation values, we raised the correlations to the power of 4.

When we analyzed the CePNEM model results, we found that only 2.7% of neurons detected as having behavioral encoding, suggesting that the vast majority (>95%) of our detected encodings are not due to overfitting.

### Constructing low-dimensional embeddings of neurons via UMAP

We wanted to use CePNEM to construct a low-dimensional UMAP space where any neuron from any animal could be embedded. To accomplish this, we took the three modeled behaviors from 12 SWF415 animals and appended them, so as to have a wide range of possible behavioral dynamics. Then, we took 4,004 median CePNEM fits (sampled from 14 SWF415 animals) and extrapolated them over the appended behavioral data, to estimate what the neuron would have done under our model over a wide range of behaviors. We then ran UMAP on the resulting 4004 × 19200 matrix to define a two-dimensional embedding space. Finally, we projected all posterior CePNEM fits from each neuron into this UMAP space to create the point cloud shown in Figure 3A. We also projected subsets of neurons based on encoding type (Figures 3B–3F), identity (Figure 5E), and dataset (Figure S3); to do this, we simply run the same projection procedure on all posterior CePNEM fits from each neuron in the subset in question (i.e. the UMAP space was the same for all embeddings shown in the paper).

### Neural trace reconstruction using principal component analysis

To determine the number of principal components needed to reconstruct each neuron, PCA was performed first on all neurons in each dataset. Neurons without high enough SNR were excluded from the analysis. We determined the SNR cutoff based on our GFP datasets. Specifically, a given neuron needed to have signal standard deviation higher than 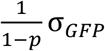, where *σ_GFB_* is the GFP signal standard deviation and *p* is the required fraction of variance explained. To reconstruct the neurons, each neuron’s loadings were sorted by absolute value. Then we increase the number of principal components used to reconstruct until the required variance explained is met. In each dataset, this process is repeated for all neurons with high enough SNR.

### Neural trace clustering analysis

To estimate the optimal number of clusters in the neural traces (Fig. S4A), we first mean center neuron. Then k-means clustering is performed on each dataset with varying number of clusters, k, ranging from 2 to 10. For each k, we compute the Calinski-Harabasz index. We repeat this on all SWF415 datasets.

### MSE model fits

For some analyses, we found it useful to fit our model in a more conventional manner, simply trying to minimize the mean-squared error (MSE) between the model fit and neural activity rather than using Gen to compute the posterior. For these fits, we deleted the noise component of our model and instead simply fit *n*[*t*] by trying to minimize the MSE between it and the observed neural activity, set *n*(0) = 0, and ignored the first 50 time points after each recording began for the MSE calculation (so for datasets with a gap in the middle, we would ignore the first 50 time points, as well as time points 801:850). We used L-BFGS for local optimization and MLSL-LDS for global optimization, and performed these fits using the NLopt Julia package (Johnson, 2022).

### Model degradation analysis

We tested how each component in the model affects the performance by quantifying the increase in error, compared to the full model, when removing the following component individually: each predictor (velocity, head curvature, feeding), the velocity non-linearity, removing the EWMA, and all non-linear features (resulting in a fully linear model). The models were fitted using our MSE fitting technique with L2 regularization. Out of the 14 pan-neuronal GCaMP baseline datasets, 5 were excluded from this analysis due to low variance in the pumping rate. 3 datasets were used to optimize the regularization parameter, and the remaining 6 datasets were used to compute the increase in error. Models were fit with 5-fold cross-validation set, splitting each dataset into 5 equal length time segments. The error was computed as the mean test time error of the cross-validation splits. For each degraded model type, neurons encoding the removed feature were selected for analysis. For example, degraded model without velocity was tested on the neurons with velocity encoding. The increase in error was computed by comparing the error in degraded model to the error of the full model. Finally, we used the Wilcoxon signed rank test for statistical significance.

### Statistical tests to examine dynamic changes in neural encoding

To determine whether a given neuron in a recording changed how it encoded behavior, we used the following procedure. First, we fit two CePNEM models to compare against each other. For baseline datasets without any stimulation (both SWF415 and NeuroPAL), we split the dataset in half and used fits from each half – the same fits used in the encoding analysis. For the heat-stimulation datasets, we took one fit from the timepoints up until just before the stimulation, and another fit from the 400 timepoint block (stim+10) to (stim+409) for heat-stimulation datasets without a gap in the middle, or alternatively (stim+10) to 800 for datasets with such a gap.

Next, we computed deconvolved activity matrices as defined above on each of the CePNEM fit posteriors. We ran the same procedure used to detect encoding, but this time instead of computing metrics on individual particles, we computed those metrics on differences between the deconvolved activity matrices for all possible pairs of particles from each of the two model fits, which was a total of slightly more than 10^8^ such differences per neuron. We used our noise threshold *ξ*_1_ as before, but *ξ*_2_ is set to 0 for this test because it is not well-defined when considering multiple model fits. Neurons that passed our encoding test at *p* = 0.05 using the differences between the deconvolved activity matrices for behaviors other than feeding (there were too few datasets with enough feeding variance in both time ranges to make a meaningful statistical comparison), and encoded behavior (using our standard behavior encoding test) in at least one time range were added to the list of encoding changing neuron candidates. Additionally, we checked whether the EWMA parameter *s* changed by computing differences between all possible values of *s* in the two model fits, and asking whether that was greater than 0. This comparison was Benjamini-Hochberg corrected over all neurons, and neurons that passed the test at *p* = 0.05 and also encoded behavior (using our standard behavior encoding test) in both time ranges were added to the list of encoding changing neuron candidates.

To additionally ensure that encoding changes are due to legitimate changes in neural relationship to behavior, and not due to model overfitting, we fit a MSE model across all timepoints and asked whether its performance was significantly worse than the performance to the two models with the hypothesized encoding difference, when evaluated over their respective time ranges. In other words, we attempted identify a model fit trained on the maximal amount of data available that explains the data as well as the two different models that we hypothesized were different. We only did this analysis for neurons in our list of encoding changing neuron candidates. The purpose of these fits was to compute the best possible explanation under the model of explaining neural activity without any assumptions about noise, so these MSE fits were not regularized at all. We then asked whether these non-regularized MSE fits could match the performance of CePNEM on both time ranges. If they could, then there exists a set of parameters that can explain neural activity in both time ranges, and so we would want to exclude the neuron in question from our encoding change analysis.

To accomplish this, for each particle in the CePNEM posterior, we extrapolated that particle into a model fit on the time range that model was trained on (excluding the first 50 time points after a recording began), and computed that particle’s MSE. Using these MSE values, we computed the probability that the CePNEM model fit’s MSE was higher than that of the conventional model evaluated over that same time range. We used Bonferroni correction on the two p-values thus computed in each of the two time ranges, and then used Benjamini-Hochberg correction across all neurons in the encoding changing neuron candidates list. The neurons that passed this test were then classified as having an encoding change.

### Encoding change strength

For neurons that had an encoding change according to our statistical framework above, it is desirable to estimate how much the neural tuning to behavior changed. To do this, we compute model fits *n*_1_ [*t*] and *n*_2_ [*t*] for each of the two time ranges in question from the median of the CePNEM posterior parameters for those time ranges. Because feeding is excluded from encoding change analysis, the feeding behavior is excluded (set to 0) for these fits. We then define encoding change strength as 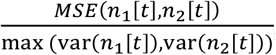, a metric of how much the neural fit changed relative to its variance. Average encoding change strength of a neuron is then the mean of its encoding change strength across each dataset where it had an encoding change.

### Behavioral analyses during cellular perturbations

For behavioral analysis in AIM- and RIC-inactivated animals, we (i) recorded animal speed on multi-worm trackers, as previously described (Rhoades et al., 2019), (ii) recorded head curvature behaviors on high-resolution single worm trackers, as previously described (Cermak et al., 2020), and (iii) quantified pharyngeal pumping manually.

---

**Supplemental Figure 1.**
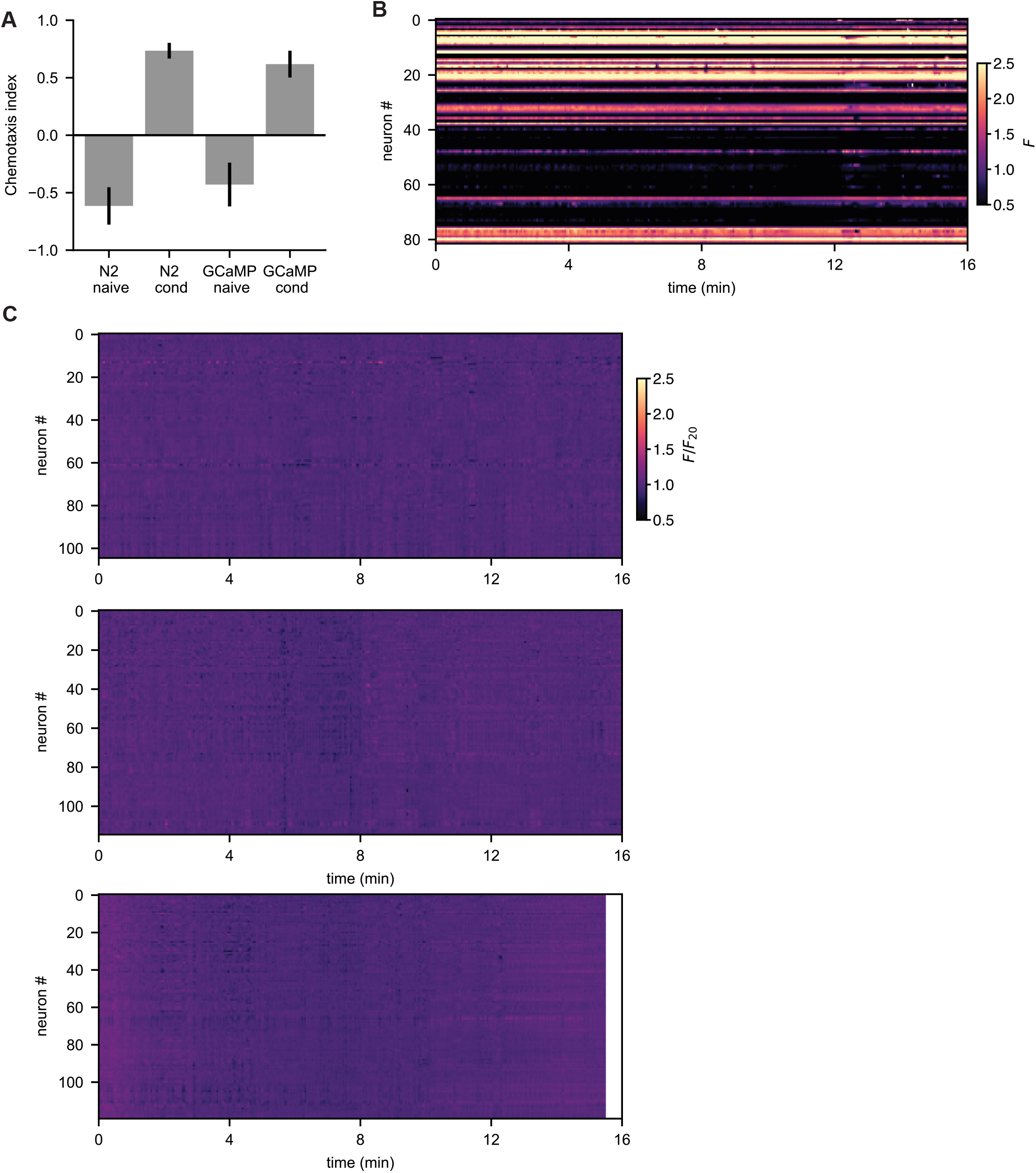
(A) Salt learning assay for N2 control animals, compared to pan-neuronal GCaMP7f animals. Naïve refers to animals grown on 0 mM NaCl; conditioned (‘cond’) refers to animals grown under the same conditions but exposed to 50mM NaCl with food for one hour prior to assay, which causes animals to prefer higher salt concentrations. Chemotaxis was measured on a plate with a 0mM to 50mM NaCl gradient with sorbitol added to ensure uniform osmolarity. Positive values correspond to chemotaxis directed toward high NaCl. Data are shown as means and standard deviation across plates. (B) Un-normalized F heatmap of neural traces collected and extracted from a control animal expressing *eat-4::NLS-GFP.* Since GFP is expressed only in a fraction of cells in this strain, perfect neural identity mapping would result in a set of bright horizontal lines (GFP-positive neurons) and a set of dark horizontal lines (GFP-negative neurons), while a registration mismatch would appear as a bright spot in the trace of an otherwise GFP-negative neuron, or a dark spot in the trace of an otherwise GFP-positive neuron. Note that there are very few instances of registration mismatches visible in the traces. As described in the main text, we estimate the number of neuron identification errors to be 0.3% of frames (see Methods). (C) F/F_20_ heatmap of neural traces collected and extracted from three GFP control animals.

**Supplemental Figure 2.**
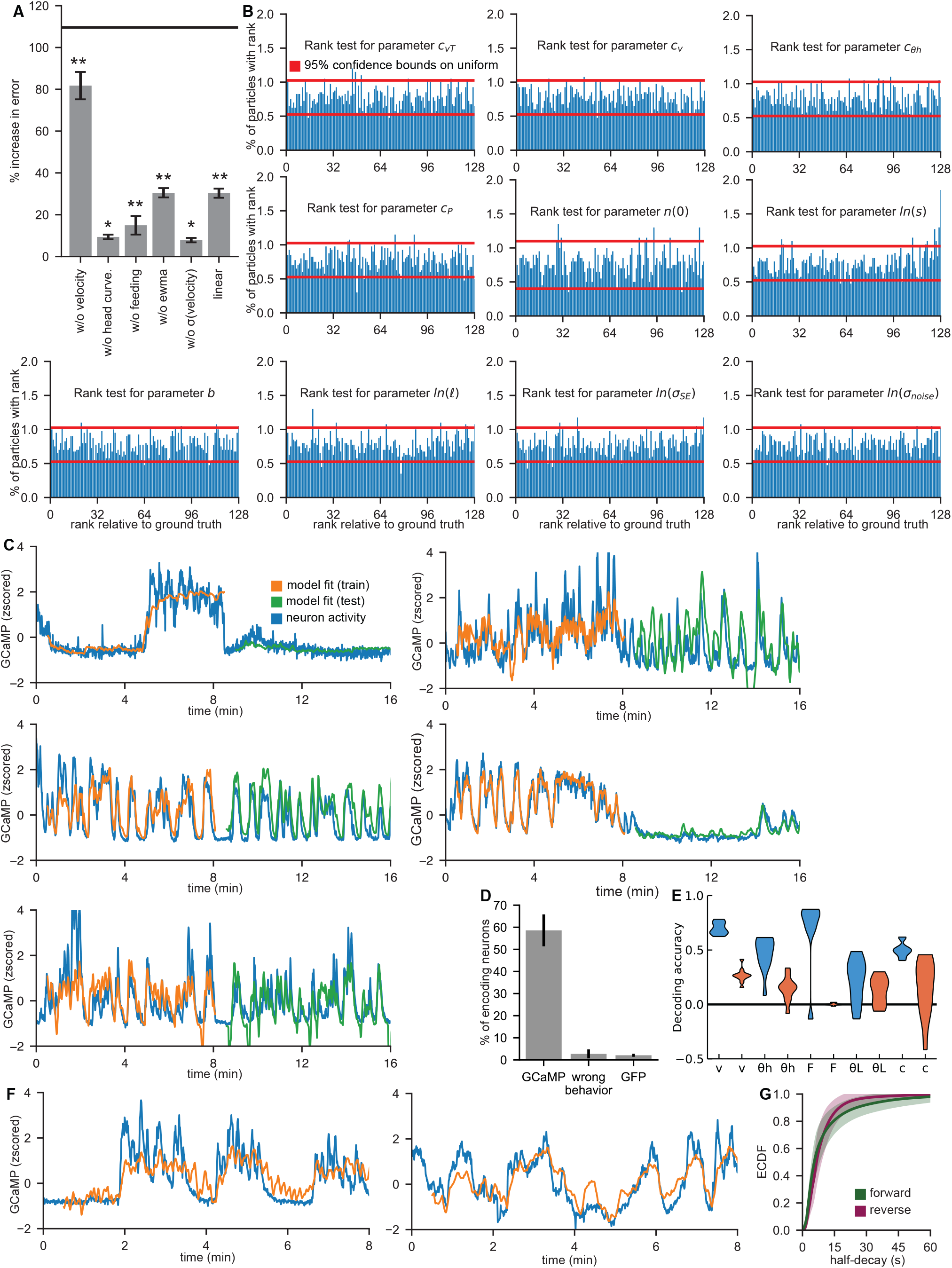
(A) Degradation analysis on each model parameter, comparing the percentage that the error (as measured by cross-validated mean-squared error when fitting the model with MSE optimization - see Methods) increases when the model is refit with that parameter removed. * = p < 0.05 ** = p < 0.0005 (Wilcoxon signed rank test). For reference, black line shows the error increase for a model with no behavioral parameters (just an offset parameter so that the model would guess each neuron’s mean activity). (B) Simulation-based calibration results for CePNEM. Simulation-based calibration was performed by simulating 1997 neurons from CePNEM using behaviors from 4 different animals and fitting them each twice, on different time ranges. For each model parameter, the ground-truth parameter was ranked within the fitted posterior. If model fitting is perfectly calibrated, the ground-truth parameter’s rank should be the uniform distribution. Therefore, for each parameter, we performed a χ^2^ test to distinguish their distribution from the uniform distribution with p=0.05. All parameters passed this test, except for the timescale parameter *s*, which has a very small calibration artifact predicted to impact <4 neurons per dataset. See Methods. (C) A series of CePNEM model fits to various neurons, showing the model’s ability to fit a wide variety of neural tunings to behavior. The model was fit on the first half of the dataset, and tested on the second half, revealing that these neurons have robust tunings to behavior across time that is well-explained by CePNEM. (D) Controls comparing the percentage of neurons that were detected as encoding behavior using real GCaMP traces with the same animal’s behavior, using the same GCaMP traces but attempting to fit with a different animal’s behavior (essentially a scramble control; ‘wrong behavior’), and using GFP datasets. See Methods. (E) Linear, L1-regularized decoder models were trained to predict various behaviors (velocity, head curvature, feeding, angular velocity, and curvature, respectively) from 11 animals from either neurons (blue) or CePNEM model residuals (orange). Decoding accuracy was assessed as 1 – MSE (decoded behavior, true behavior), averaged over five 80/20 cross-validation splits (see Methods). Note that the decoder models do much worse when trained on CePNEM model residuals than when trained on the full neural data, suggesting that the model can explain most neural variance overtly related to behavior. (F) Two neurons that encode angular velocity (defined as longer-timescale head curvature; due to the higher frequency nature of head curvature oscillations, longer-timescale is defined here as at least 5 seconds). Left: this neuron has a half-decay of *τ*_1/2_ = 10.0 ± 2.8 seconds, while the neuron on the right has a half-decay of *τ*_1/2_ = 9.5 ± 1.3 seconds. The neuron on the right is multiplexed with velocity as well. (G) Mean ECDF of the model half-decay time of all neurons demonstrated to encode forward locomotion, contrasted with the ECDF of neurons demonstrated to encode reverse locomotion, in 14 animals. The shaded regions represent the standard deviation between animals. The median fraction (across animals) of forward neurons with long timescales (half-decay *τ*_1/2_ > 20s) was 0.12, compared with only 0.03 for reversal neurons; this difference was statistically significant (p = 0.029) under a Mann-Whitney U-Test.

**Supplemental Figure 3.**
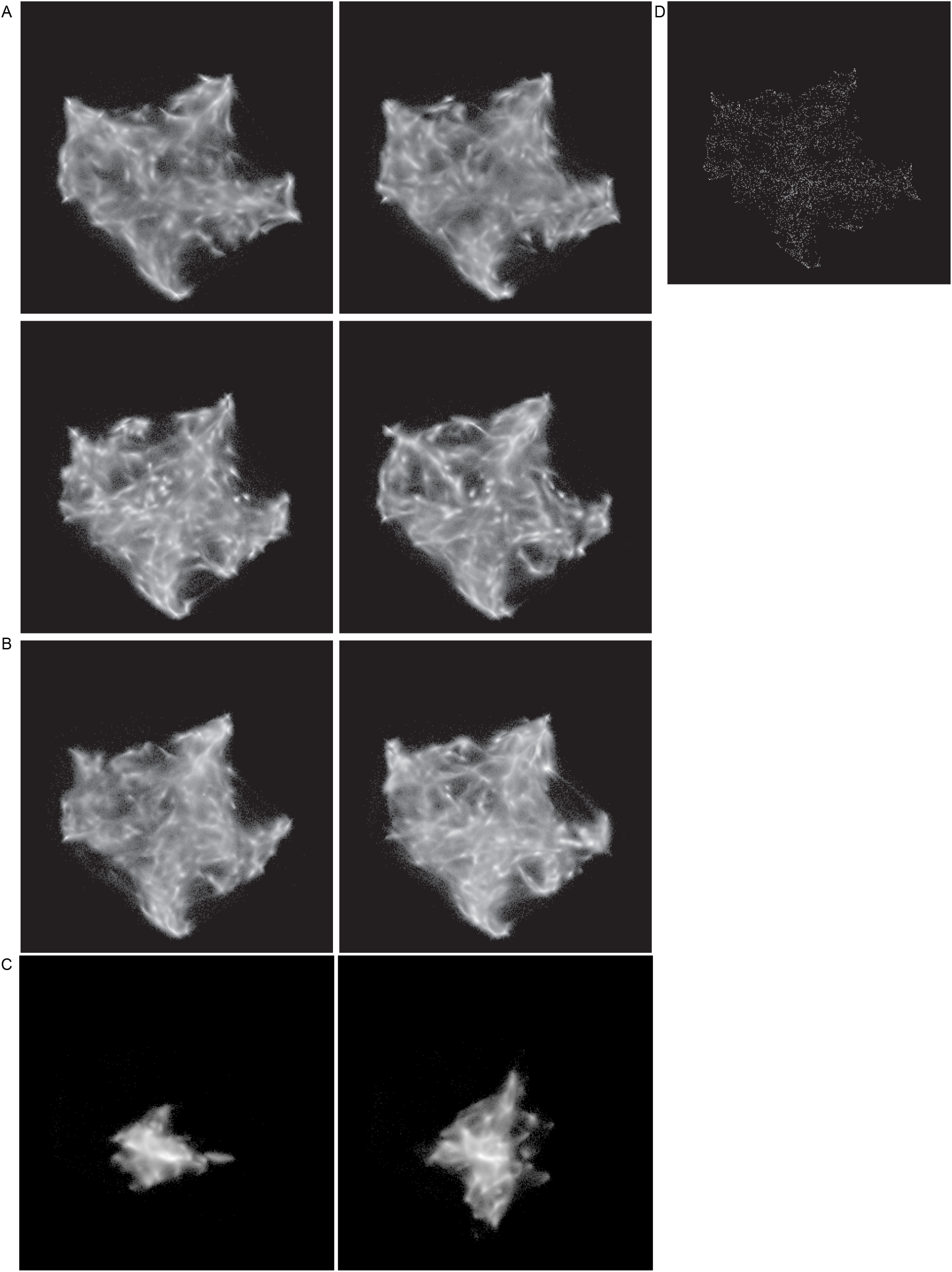
(A) Projections of all neurons from each of four different SWF415 animals into the same UMAP space (built from full population of animals; same as in Fig. 3A). Observe that the overall structure is very similar, suggesting that the locations of neurons in UMAP space are similar across datasets. (B) Projections of all neurons from each of two different NeuroPAL animals into the UMAP space. These neurons also fill in a similar pattern to that of the SWF415 animals, suggesting that the overall neural encodings of the two strains are similar. (C) Projections of all neurons from each of two different GFP control animals into the UMAP space. These neurons fail to fill most of the space, which is consistent with the non-encoding nature of neurons in this control strain. (D) Projections of all neurons from 14 different SWF415 animals into the UMAP space, taking the median of each neuron’s posterior point cloud in the UMAP space. Note that the medians fill out the same space as when projecting the full posteriors, suggesting the continuity of the UMAP space is not merely an artifact of parameter uncertainty.

**Supplemental Figure 4.**
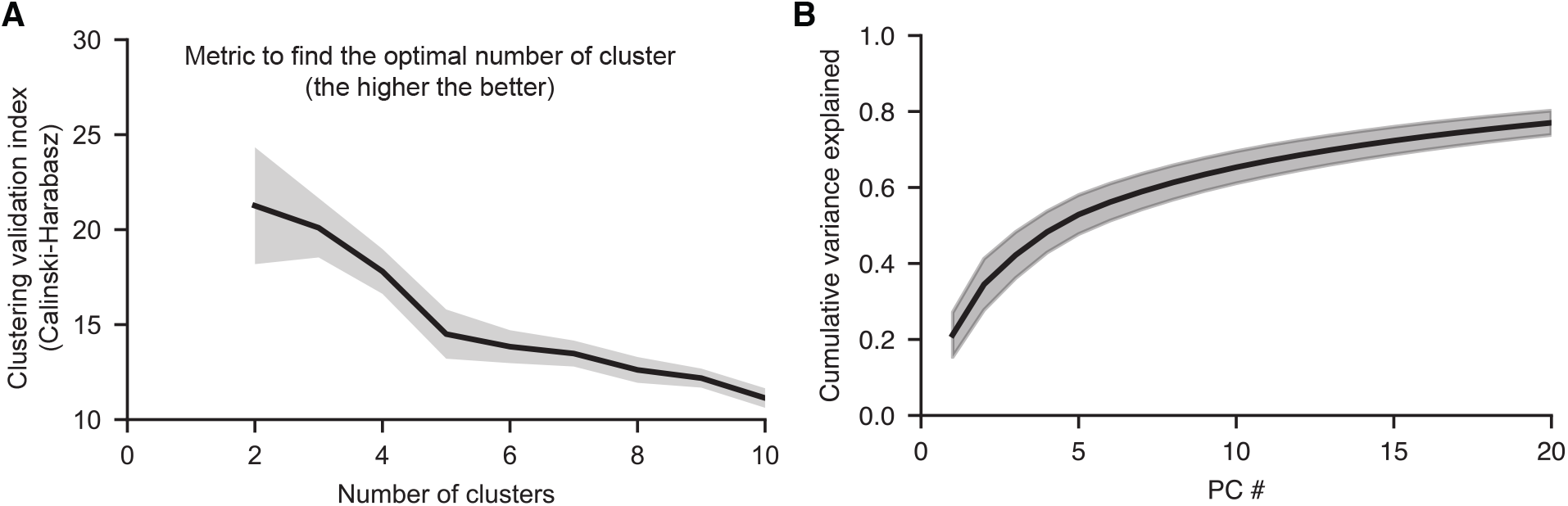
(A) An analysis of clusterability of all neurons that encode behavior. For each dataset, we attempted to cluster all neurons that encode behavior using a similarity metric based on the difference of the neurons’ GCaMP traces. To determine the optimal number of clusters, we computed the Calinski-Harabasz index over varying number of clusters when performing k-means clustering on the neural traces. Clustering was done on a per dataset basis on all SWF415 datasets, and the mean and standard error values are plotted. Note that the optimal number of clusters in this analysis is 2, which is the minimum number that can be assessed with this metric. This suggests that there is not a larger set of discrete subgroups of neurons that are separable from one another. (B) Cumulative variance explained by the top 20 PCs, averaged over 14 animals. The shaded region is the standard deviation across animals.

**Supplemental Figure 5.**
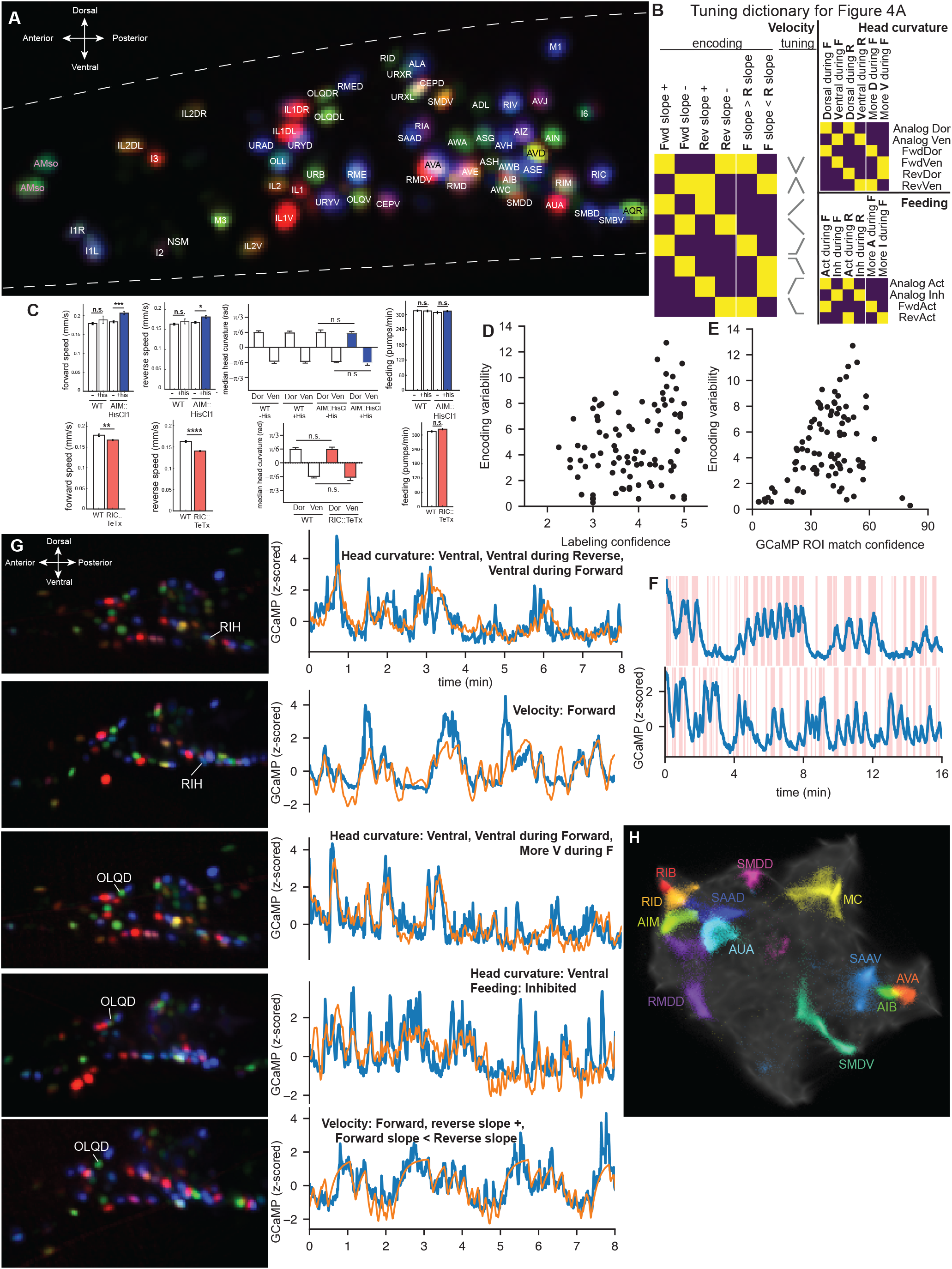
(A) A RGB composite image of one of the NeuroPAL animals that we recorded. The composite was constructed by combining images of NLS-mTagBFP2 (shown in blue), NLS-cyOFP2 (shown in green), and NLS-mNeptune2.5 (shown in red). Using this composite image, we were able to label a large number of neurons in this animal. Neural identity was determined while making use of all 3D information, but for display purposes here we show a maximum intensity projection of a subset of the z-slices from the recording. Therefore, this image does not show all the neurons in the head (a maximum intensity projection of all z-slices is too dense with neurons to show for display purposes here). (B) A legend that demonstrates how to link the encodings in Figure 4A with the neuron tunings in Figures 2E-G. To perform this mapping, first choose the behavior (velocity, head curvature, or feeding) and reference the corresponding section of the legend. Then, look at the last six columns of Figure 4A of the chosen behavior for the neuron in question (which correspond to the six columns shown here). Match the pattern shown in those columns with one of the rows of the miniature heatmap shown here (if you cannot find such a match, it is likely that there was insufficient statistical power to ascribe a tuning to that neuron). Finally, read the tuning corresponding to that row; these tunings will be in the same form as in Figures 2E-G (for velocity, this will take the form of a tuning diagram, and for head curvature and feeding it will take the form of a tuning category). (C) Effects of perturbing AIM and RIC neurons on the animal’s behavioral output. AIM was inactivated via chemogenetic silencing using the Histamine-gated chloride channel (HisCl) and RIC was inactivated via tetanus toxin expression. For both perturbations, we quantified forward speed, reverse speed, median head curvature during dorsal and ventral head bends, and feeding rates. *p<0.05, **p<0.01, ***p<0.001, ****p<0.0001, Bonferroni-corrected t-test. n.s., not significant. (D) Scatter plot of labeling confidence (a qualitative metric determined by person scoring, reflecting their confidence that the neuron is correctly identified based on position and multi-spectral fluorescence; the higher the better; note that neurons with sufficiently low confidence were entirely excluded from all analyses in the paper, and this plot only shows values above this threshold) and encoding variability (lower value means more consistency). There is no evident relationship between these values, suggesting that labeling error does not introduce encoding variability. (E) Scatter plot of GCaMP ROI match score (the higher the better; see Methods) and encoding variability shows no relationship. This suggests that the process that matches the NeuroPAL ROI to the GCaMP ROI does not introduce encoding variability. (F) Example traces of neuron AVA from 2 different animals to show the previously-described reliable tuning to reversals. Red shading indicates reversals. (G) Examples of the variable coupling neurons (RIH from 2 animals and OLQD from 3 animals shown). On the left column, the NeuroPAL fluorescence images with the neurons identified show consistent color combination and location within a given neuron class. On the right column, the corresponding neural traces (blue) are shown along with CePNEM fits (orange), and a written description of the encoding properties. Note that the neurons of the same class from different animals encode different sets of behaviors. (H) UMAP plot showing the posterior distributions of the CePNEM model fits for various neurons; each neuron is plotted in a different color. The same set of time points from the same animal were used for each neuron’s fit. This plot shows a subset of neurons with largely nonoverlapping tunings, just to illustrate how neurons map onto the UMAP space described in Fig. 3.

**Supplemental Figure 6.**
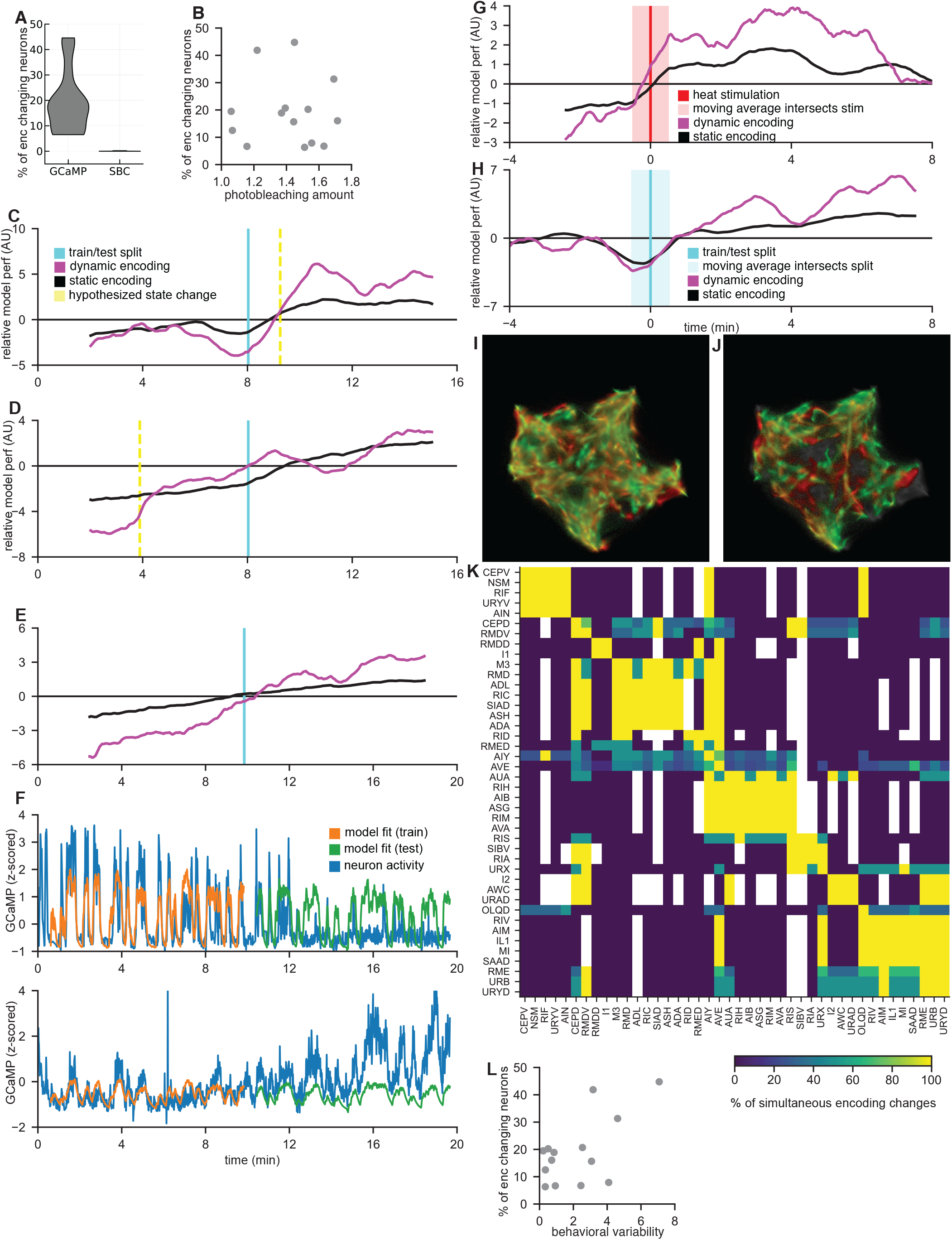
(A) An analysis of what fraction of neurons were detected as changing encoding in our GCaMP datasets and simulated datasets. Simulated datasets are labeled ‘SBC’ for simulation-based calibrations. These are neurons simulated from the CePNEM model, where ground-truth parameters were set to not have any encoding changes. (B) Scatterplot of datasets showing that extent of photobleaching is not correlated with detection of encoding changes. (C) The same dataset in Fig. 5A but also plotting the relative model performance averaged over the static encoding neurons. Note that the black line does not show the sudden changes in value seen for the purple line. (D) Same as (C), but for the dataset in Fig. 5C. (E) An example dataset that shows a less synchronized encoding change. (F) Two example encoding changing neurons from the animal in (E), one with an abrupt encoding change at approximately 12 minutes, and another neuron that appears to have a slowly-increasing gain to its behavioral encoding over the last ~10 minutes of the recording. (G-H) A comparison of the relative model performance averaged across all 11 animals that underwent a heat shock (G; same as Fig. 4G) with the same metric computed over 4 animals that were not stimulated (H). Note that the baseline animals do not have a sharp change in relative model performance at the train/test split, suggesting that the encoding changes in the heatstimulation datasets are a direct result of the stimulation. (I) UMAP of all encoding changing neurons in non-heatstim SWF415 animals. The projections of all neurons in the first time segment (before their encoding change) are shown in red; the projections of all neurons in the second time segment (after their encoding change) are shown in green. Observe that neurons throughout encoding space can exhibit encoding changes. (J) The same analysis as (I), except these animals were subjected to a heat stimulation, and the encoding changes are measured before vs after the stimulus. (K) A matrix representing which pairs of neurons have simultaneous encoding changes. Each row represents the probability that the neuron corresponding to that row had an encoding change conditioned on the column neuron changing its encoding. This probability was computed only over the set of datasets where both neurons were detected; if this set was empty, the corresponding entry in the matrix is left blank (white). The matrix was hierarchically clustered, and exhibits striking stereotypy. (L) A plot of the fraction of encoding neurons that exhibited encoding change in a dataset, compared with the behavioral difference between the first and second half of that dataset. Behavioral variability was computed as the sum of the absolute values of the differences (across the two time segments) of the following behavioral parameters (each such parameter was normalized to the standard deviation of that behavior across all 14 SWF415 datasets): median of reverse velocity, median of forward velocity, 25^th^ percentile of head curvature, 75^th^ percentile of head curvature, 25^th^ percentile of feeding rate, and 75^th^ percentile of feeding rate. This value provides a general description of how much the distributions of behavioral parameters changed across the two halves of the recording. Observe that datasets with large behavioral changes tend to have more encoding changes, suggesting that the neural flexibility may be related to the observed behavior changing.

## Supplemental Item Legends

**Movie S1. Example video of baseline recording conditions.** A two minute-long excerpt from an example neural/behavioral dataset, showing the NIR behavioral recording. Raw video data is shown with overlaid information: (i) blue, orange, and green dots are the identified targets for worm tracking that were determined during live recording, which allowed us to locate the worm’s head and keep the animal centered in view; (ii) black line shows a spline fit to the animal’s centerline; (iii) upper left shows time and values of three ongoing behavioral parameters: velocity, head curvature, and feeding rate.

## Notes

### Competing Interest Statement

The authors have declared no competing interest.

## REFERENCES

Allen, W.E., Chen, M.Z., Pichamoorthy, N., Tien, R.H., Pachitariu, M., Luo, L., Deisseroth, K., 2019. Thirst regulates motivated behavior through modulation of brainwide neural population dynamics. Science 364, 253. https://doi.org/10.1126/science.aav3932

Anderson, D.J., Adolphs, R., 2014. A framework for studying emotions across species. Cell 157, 187–200. https://doi.org/10.1016/j.cell.2014.03.003

Ardiel, E.L., Yu, A.J., Giles, A.C., Rankin, C.H., 2017. Habituation as an adaptive shift in response strategy mediated by neuropeptides. NPJ Sci Learn 2, 9. https://doi.org/10.1038/s41539-017-0011-8

Berzuini, C., Gilks, W., 2001. RESAMPLE-MOVE Filtering with Cross-Model Jumps, in: Doucet, A., de Freitas, N., Gordon, N. (Eds.), Sequential Monte Carlo Methods in Practice, Statistics for Engineering and Information Science. Springer, New York, NY, pp. 117–138. https://doi.org/10.1007/978-1-4757-3437-9_6

Brezovec, L.E., Berger, A.B., Druckmann, S., Clandinin, T.R., 2022. Mapping the Neural Dynamics of Locomotion across the Drosophila Brain. bioRxiv. https://doi.org/10.1101/2022.03.20.485047

Byrne Rodgers, J., Ryu, W.S., 2020. Targeted thermal stimulation and high-content phenotyping reveal that the C. elegans escape response integrates current behavioral state and past experience. PLoS One 15, e0229399. https://doi.org/10.1371/journal.pone.0229399

Cermak, N., Yu, S.K., Clark, R., Huang, Y.-C., Baskoylu, S.N., Flavell, S.W., 2020. Whole-organism behavioral profiling reveals a role for dopamine in state-dependent motor program coupling in C. elegans. Elife 9, e57093. https://doi.org/10.7554/eLife.57093

Chalfie, M., Sulston, J.E., White, J.G., Southgate, E., Thomson, J.N., Brenner, S., 1985. The neural circuit for touch sensitivity in Caenorhabditis elegans. J Neurosci 5, 956–964.

Chew, Y.L., Tanizawa, Y., Cho, Y., Zhao, B., Yu, A.J., Ardiel, E.L., Rabinowitch, I., Bai, J., Rankin, C.H., Lu, H., Beets, I., Schafer, W.R., 2018. An Afferent Neuropeptide System Transmits Mechanosensory Signals Triggering Sensitization and Arousal in C. elegans. Neuron 99, 1233–1246.e6. https://doi.org/10.1016/j.neuron.2018.08.003

Cook, S.J., Jarrell, T.A., Brittin, C.A., Wang, Y., Bloniarz, A.E., Yakovlev, M.A., Nguyen, K.C.Q., Tang, L.T.-H., Bayer, E.A., Duerr, J.S., Bülow, H.E., Hobert, O., Hall, D.H., Emmons, S.W., 2019. Whole-animal connectomes of both Caenorhabditis elegans sexes. Nature 571, 63–71. https://doi.org/10.1038/s41586-019-1352-7

Croll, N.A., 1975. Components and patterns in the behaviour of the nematode Caenorhabditis elegans. Journal of Zoology 176, 159–176. https://doi.org/10.1111/j.1469-7998.1975.tb03191.x

Cusumano-Towner, M.F., Saad, F.A., Lew, A.K., Mansinghka, V.K., 2019. Gen: a general-purpose probabilistic programming system with programmable inference, in: Proceedings of the 40th ACM SIGPLAN Conference on Programming Language Design and Implementation, PLDI 2019. Association for Computing Machinery, New York, NY, USA, pp. 221–236. https://doi.org/10.1145/3314221.3314642

Dana, H., Sun, Y., Mohar, B., Hulse, B.K., Kerlin, A.M., Hasseman, J.P., Tsegaye, G., Tsang, A., Wong, A., Patel, R., Macklin, J.J., Chen, Y., Konnerth, A., Jayaraman, V., Looger, L.L., Schreiter, E.R., Svoboda, K., Kim, D.S., 2019. High-performance calcium sensors for imaging activity in neuronal populations and microcompartments. Nat Methods 16, 649–657. https://doi.org/10.1038/s41592-019-0435-6

Flavell, S.W., Gogolla, N., Lovett-Barron, M., Zelikowsky, M., 2022. The emergence and influence of internal states. Neuron 110, 2545–2570. https://doi.org/10.1016/j.neuron.2022.04.030

Flavell, S.W., Pokala, N., Macosko, E.Z., Albrecht, D.R., Larsch, J., Bargmann, C.I., 2013. Serotonin and the neuropeptide PDF initiate and extend opposing behavioral states in C. elegans. Cell 154, 1023–1035. https://doi.org/10.1016/j.cell.2013.08.001

Flavell, S.W., Raizen, D.M., You, Y.-J., 2020. Behavioral States. Genetics 216, 315–332. https://doi.org/10.1534/genetics.120.303539

Fujiwara, M., Sengupta, P., McIntire, S.L., 2002. Regulation of body size and behavioral state of C. elegans by sensory perception and the EGL-4 cGMP-dependent protein kinase. Neuron 36, 1091–1102.

Gordus, A., Pokala, N., Levy, S., Flavell, S.W., Bargmann, C.I., 2015. Feedback from network states generates variability in a probabilistic olfactory circuit. Cell 161, 215–227. https://doi.org/10.1016/j.cell.2015.02.018

Guizar-Sicairos, M., Thurman, S.T., Fienup, J.R., 2008. Efficient subpixel image registration algorithms. Opt Lett 33, 156–158. https://doi.org/10.1364/ol.33.000156

Hallinen, K.M., Dempsey, R., Scholz, M., Yu, X., Linder, A., Randi, F., Sharma, A.K., Shaevitz, J.W., Leifer, A.M., 2021. Decoding locomotion from population neural activity in moving C. elegans. Elife 10, e66135. https://doi.org/10.7554/eLife.66135

Hendricks, M., Ha, H., Maffey, N., Zhang, Y., 2012. Compartmentalized calcium dynamics in a C. elegans interneuron encode head movement. Nature 487, 99–103. https://doi.org/10.1038/nature11081

Ji, N., Madan, G.K., Fabre, G.I., Dayan, A., Baker, C.M., Kramer, T.S., Nwabudike, I., Flavell, S.W., 2021. A neural circuit for flexible control of persistent behavioral states. Elife 10, e62889. https://doi.org/10.7554/eLife.62889

Johnson, S.G., 2022. stevengj/nlopt.

Kaplan, H.S., Salazar Thula, O., Khoss, N., Zimmer, M., 2020. Nested Neuronal Dynamics Orchestrate a Behavioral Hierarchy across Timescales. Neuron 105, 562–576.e9. https://doi.org/10.1016/j.neuron.2019.10.037

Kato, S., Kaplan, H.S., Schrödel, T., Skora, S., Lindsay, T.H., Yemini, E., Lockery, S., Zimmer, M., 2015. Global brain dynamics embed the motor command sequence of Caenorhabditis elegans. Cell 163, 656–669. https://doi.org/10.1016/j.cell.2015.09.034

Kotera, I., Tran, N.A., Fu, D., Kim, J.H., Byrne Rodgers, J., Ryu, W.S., 2016. Pan-neuronal screening in Caenorhabditis elegans reveals asymmetric dynamics of AWC neurons is critical for thermal avoidance behavior. Elife 5, e19021. https://doi.org/10.7554/eLife.19021

Li, Z., Liu, J., Zheng, M., Xu, X.Z.S., 2014. Encoding of both analog- and digital-like behavioral outputs by one C. elegans interneuron. Cell 159, 751–765. https://doi.org/10.1016/j.cell.2014.09.056

Lim, M.A., Chitturi, J., Laskova, V., Meng, J., Findeis, D., Wiekenberg, A., Mulcahy, B., Luo, L., Li, Y., Lu, Y., Hung, W., Qu, Y., Ho, C.-Y., Holmyard, D., Ji, N., McWhirter, R., Samuel, A.D., Miller, D.M., Schnabel, R., Calarco, J.A., Zhen, M., 2016. Neuroendocrine modulation sustains the C. elegans forward motor state. Elife 5, e19887. https://doi.org/10.7554/eLife.19887

Luo, L., Wen, Q., Ren, J., Hendricks, M., Gershow, M., Qin, Y., Greenwood, J., Soucy, E.R., Klein, M., Smith-Parker, H.K., Calvo, A.C., Colón-Ramos, D.A., Samuel, A.D.T., Zhang, Y., 2014. Dynamic encoding of perception, memory, and movement in a C. elegans chemotaxis circuit. Neuron 82, 1115–1128. https://doi.org/10.1016/j.neuron.2014.05.010

Marques, J.C., Li, M., Schaak, D., Robson, D.N., Li, J.M., 2020. Internal state dynamics shape brainwide activity and foraging behaviour. Nature 577, 239–243. https://doi.org/10.1038/s41586-019-1858-z

Mathis, A., Mamidanna, P., Cury, K.M., Abe, T., Murthy, V.N., Mathis, M.W., Bethge, M., 2018. DeepLabCut: markerless pose estimation of user-defined body parts with deep learning. Nat Neurosci 21, 1281–1289. https://doi.org/10.1038/s41593-018-0209-y

Musall, S., Kaufman, M.T., Juavinett, A.L., Gluf, S., Churchland, A.K., 2019. Single-trial neural dynamics are dominated by richly varied movements. Nat Neurosci 22, 1677–1686. https://doi.org/10.1038/s41593-019-0502-4

Neal, R.M., 2011. MCMC using Hamiltonian dynamics. https://doi.org/10.1201/b10905

Nguyen, J.P., Shipley, F.B., Linder, A.N., Plummer, G.S., Liu, M., Setru, S.U., Shaevitz, J.W., Leifer, A.M., 2016. Whole-brain calcium imaging with cellular resolution in freely behaving Caenorhabditis elegans. Proc. Natl. Acad. Sci. U.S.A. 113, E1074–1081. https://doi.org/10.1073/pnas.1507110112

Niell, C.M., Stryker, M.P., 2010. Modulation of visual responses by behavioral state in mouse visual cortex. Neuron 65, 472–479. https://doi.org/10.1016/j.neuron.2010.01.033

Raizen, D.M., Zimmerman, J.E., Maycock, M.H., Ta, U.D., You, Y., Sundaram, M.V., Pack, A.I., 2008. Lethargus is a Caenorhabditis elegans sleep-like state. Nature 451, 569–572. https://doi.org/10.1038/nature06535

Rhoades, J.L., Nelson, J.C., Nwabudike, I., Yu, S.K., McLachlan, I.G., Madan, G.K., Abebe, E., Powers, J.R., Colón-Ramos, D.A., Flavell, S.W., 2019. ASICs Mediate Food Responses in an Enteric Serotonergic Neuron that Controls Foraging Behaviors. Cell 176, 85–97.e14. https://doi.org/10.1016/j.cell.2018.11.023

Roberts, W.M., Augustine, S.B., Lawton, K.J., Lindsay, T.H., Thiele, T.R., Izquierdo, E.J., Faumont, S., Lindsay, R.A., Britton, M.C., Pokala, N., Bargmann, C.I., Lockery, S.R., 2016. A stochastic neuronal model predicts random search behaviors at multiple spatial scales in C. elegans. Elife 5, e12572. https://doi.org/10.7554/eLife.12572

Rudin, L.I., Osher, S., Fatemi, E., 1992. Nonlinear total variation based noise removal algorithms. Physica D: Nonlinear Phenomena 60, 259–268. https://doi.org/10.1016/0167-2789(92)90242-F

Schaffer, E.S., Mishra, N., Whiteway, M.R., Li, W., Vancura, M.B., Freedman, J., Patel, K.B., Voleti, V., Paninski, L., Hillman, E.M.C., Abbott, L.F., Axel, R., 2021. Flygenvectors: The spatial and temporal structure of neural activity across the fly brain. bioRxiv. https://doi.org/10.1101/2021.09.25.461804

Stringer, C., Pachitariu, M., Steinmetz, N., Reddy, C.B., Carandini, M., Harris, K.D., 2019. Spontaneous behaviors drive multidimensional, brainwide activity. Science 364, 255. https://doi.org/10.1126/science.aav7893

Talts, S., Betancourt, M., Simpson, D., Vehtari, A., Gelman, A., 2020. Validating Bayesian Inference Algorithms with Simulation-Based Calibration. https://doi.org/10.48550/arXiv.1804.06788

Urai, A.E., Doiron, B., Leifer, A.M., Churchland, A.K., 2022. Large-scale neural recordings call for new insights to link brain and behavior. Nat Neurosci 25, 11–19. https://doi.org/10.1038/s41593-021-00980-9

Van Buskirk, C., Sternberg, P.W., 2007. Epidermal growth factor signaling induces behavioral quiescence in Caenorhabditis elegans. Nat Neurosci 10, 1300–1307. https://doi.org/10.1038/nn1981

Venkatachalam, V., Ji, N., Wang, X., Clark, C., Mitchell, J.K., Klein, M., Tabone, C.J., Florman, J., Ji, H., Greenwood, J., Chisholm, A.D., Srinivasan, J., Alkema, M., Zhen, M., Samuel, A.D.T., 2016. Pan-neuronal imaging in roaming Caenorhabditis elegans. Proc. Natl. Acad. Sci. U.S.A. 113, E1082–1088. https://doi.org/10.1073/pnas.1507109113

Wang, Y.L., Jaklitsch, E.L., Grooms, N.W.F., Schulting, L.G., Chung, S.H., 2022. High-throughput submicron-resolution microscopy of entire C. elegans populations under strong immobilization by cooling cultivation plates. https://doi.org/10.1101/2021.12.09.471981

Wei, Z., Lin, B.-J., Chen, T.-W., Daie, K., Svoboda, K., Druckmann, S., 2020. A comparison of neuronal population dynamics measured with calcium imaging and electrophysiology. PLoS Comput Biol 16, e1008198. https://doi.org/10.1371/journal.pcbi.1008198

White, J.G., Southgate, E., Thomson, J.N., Brenner, S., 1986. The structure of the nervous system of the nematode Caenorhabditis elegans. Philos. Trans. R. Soc. Lond., B, Biol. Sci. 314, 1–340.

Witvliet, D., Mulcahy, B., Mitchell, J.K., Meirovitch, Y., Berger, D.R., Wu, Y., Liu, Y., Koh, W.X., Parvathala, R., Holmyard, D., Schalek, R.L., Shavit, N., Chisholm, A.D., Lichtman, J.W., Samuel, A.D.T., Zhen, M., 2021. Connectomes across development reveal principles of brain maturation. Nature 596, 257–261. https://doi.org/10.1038/s41586-021-03778-8

Wolny, A., Cerrone, L., Vijayan, A., Tofanelli, R., Barro, A.V., Louveaux, M., Wenzl, C., Strauss, S., Wilson-Sánchez, D., Lymbouridou, R., Steigleder, S.S., Pape, C., Bailoni, A., Duran-Nebreda, S., Bassel, G.W., Lohmann, J.U., Tsiantis, M., Hamprecht, F.A., Schneitz, K., Maizel, A., Kreshuk, A., 2020. Accurate and versatile 3D segmentation of plant tissues at cellular resolution. Elife 9, e57613. https://doi.org/10.7554/eLife.57613

Yemini, E., Lin, A., Nejatbakhsh, A., Varol, E., Sun, R., Mena, G.E., Samuel, A.D.T., Paninski, L., Venkatachalam, V., Hobert, O., 2021. NeuroPAL: A Multicolor Atlas for Whole-Brain Neuronal Identification in C. elegans. Cell 184, 272–288.e11. https://doi.org/10.1016/j.cell.2020.12.012

Zhang, M., Chung, S.H., Fang-Yen, C., Craig, C., Kerr, R.A., Suzuki, H., Samuel, A.D.T., Mazur, E., Schafer, W.R., 2008. A self-regulating feed-forward circuit controlling C. elegans egg-laying behavior. Curr Biol 18, 1445–1455. https://doi.org/10.1016/j.cub.2008.08.047

